# Functional diversity of a temperate continental flora: Wide but skewed coverage of the global spectrum of plant form and function

**DOI:** 10.1101/2025.07.10.664146

**Authors:** Eugenia Sánchez-Díaz, Sandra Díaz

## Abstract

1. Habitat filtering is assumed to promote plant functional convergence, but local functional ranges are often wide. Using the Global Spectrum of Plant Form and Function (GSPFF) as global reference, we tested whether a regional flora shared the same trait trade-offs as the global flora, and whether the regional phenospace supports the habitat filtering hypothesis.
2. We measured six key functional traits in 268 central Argentina species. We compared this flora with the GSPFF using univariate and multivariate methods. We tested whether the regional phenospace was smaller or more skewed than expected by chance, and analysed its ‘topography’.
3. Regional and global phenospaces shared major axes, syndromes, and trait correlations. Despite having fewer species, the regional flora broadly covered the GSPFF, but without tall, large-leaved, large-seeded trees. The most frequent species were conservative herbs and shrubs. Distinct species also showed highly conservative attributes.
4. Despite being moderate to harsh, central Argentina environmental conditions do not appear to preclude the regional presence of most functional syndromes previously identified at the global scale. However, the regional and global floras differ in the most frequent syndromes. At the regional scale, habitat filtering shapes functional ‘topographies’ more than it constrains the extent of phenospaces.

## 1 INTRODUCTION

The taxonomic and functional composition of communities is the result of a complex interplay of biogeographical factors, such as landscape connectivity and evolutionary history, along with abiotic and biotic filters (Díaz et al., 1998a; Keddy, 1992; Lomolino et al., 2017; Violle et al., 2007; Weiher et al., 1998). Acting on the species pool, these filters shape species assemblages and lead to functional convergence, i.e. a reduction in the range of growth forms or trait syndromes present, or the consistent predominance of particular ones, under specific environmental conditions (Cornwell et al., 2006; de Bello et al., 2009; Díaz et al., 1998b). Functional convergence driven by habitat filtering lies at the core of phytogeography. While its roots are ancient –traceable to Humboldt’s observations of plant life form changes across altitudinal gradients (Von Humboldt & Bonpland, 1807)– the concept continues to be relevant today (Bruelheide et al., 2018; Swenson et al., 2012). However, reports of convergence coexist with an apparently contradictory observation: at any given local site, the range of viable plant functional syndromes is remarkably wide (Westoby et al., 2002), often explained by microsite heterogeneity or partition of resources (functional divergence of coexisting species, Cornwell & Ackerly, 2009).

Functional attributes are a crucial factor influencing species inclusion in, or exclusion from, the assemblage of a given regional landscape or local community (Moles et al., 2009, 2014; Shipley et al., 2006; Swenson et al., 2012), but the empirical testing of this influence has lagged behind. This gap is partly due to the difficulty in finding an adequate global functional space of reference or ‘functional pool’ with which to compare more restricted species sets. However, with the publication of the Global Spectrum of Plant Form and Function (hereafter GSPFF; Díaz et al., 2016) we now have an effective functional reference to compare regional or local plant assemblages. The GSPFF describes the functional space occupied by vascular plants worldwide in six traits crucial to growth, survival, and reproduction. It encompasses species from most of the major climatic and geographic regions (Fig. 1 and Extended Data Figure 1 of Díaz et al., 2016), with herbaceous non-graminoid plants comprising 33% of the database, trees 27%, shrubs 15%, herbaceous graminoids 9%, and the remaining 16% including climbers, succulents, ferns, and other growth forms (Díaz et al., 2022). Almost three quarters of the trait variation contained in the GSPFF is captured by two multivariate dimensions, one reflecting the size of whole plants and their parts (going from short plants with small seeds to tall species with large seeds), and the other reflecting the leaf economics spectrum (from plants with ‘acquisitive’ leaves, which are cheaply constructed and with short expected lifespans, to ‘conservative’ leaves, which are high-cost and have longer expected lifespans and higher expected survival to abiotic and biotic hazards). The GSPFF, in the authors’ words, provides “a backdrop against which many other facets of plant biology can be placed into a broader context [because] trait variation in any given physical setting can be compared to the worldwide background.” So far, the GSPFF has been used as a reference to compare a small number of regional or local floras, expected to be highly functionally constrained because of climatic harshness (e.g., Thomas et al., 2020) or insularity (e.g., Barajas Barbosa et al., 2023). Noteworthy, one limitation to the comparison of local or regional floras to the GSPFF is that, despite the explosive development of functional trait databases, there is still a limited number of species worldwide with available empirical trait measurements across different organs, which are necessary to characterize trait syndromes (Kattge et al., 2020).

**FIGURE 1.**
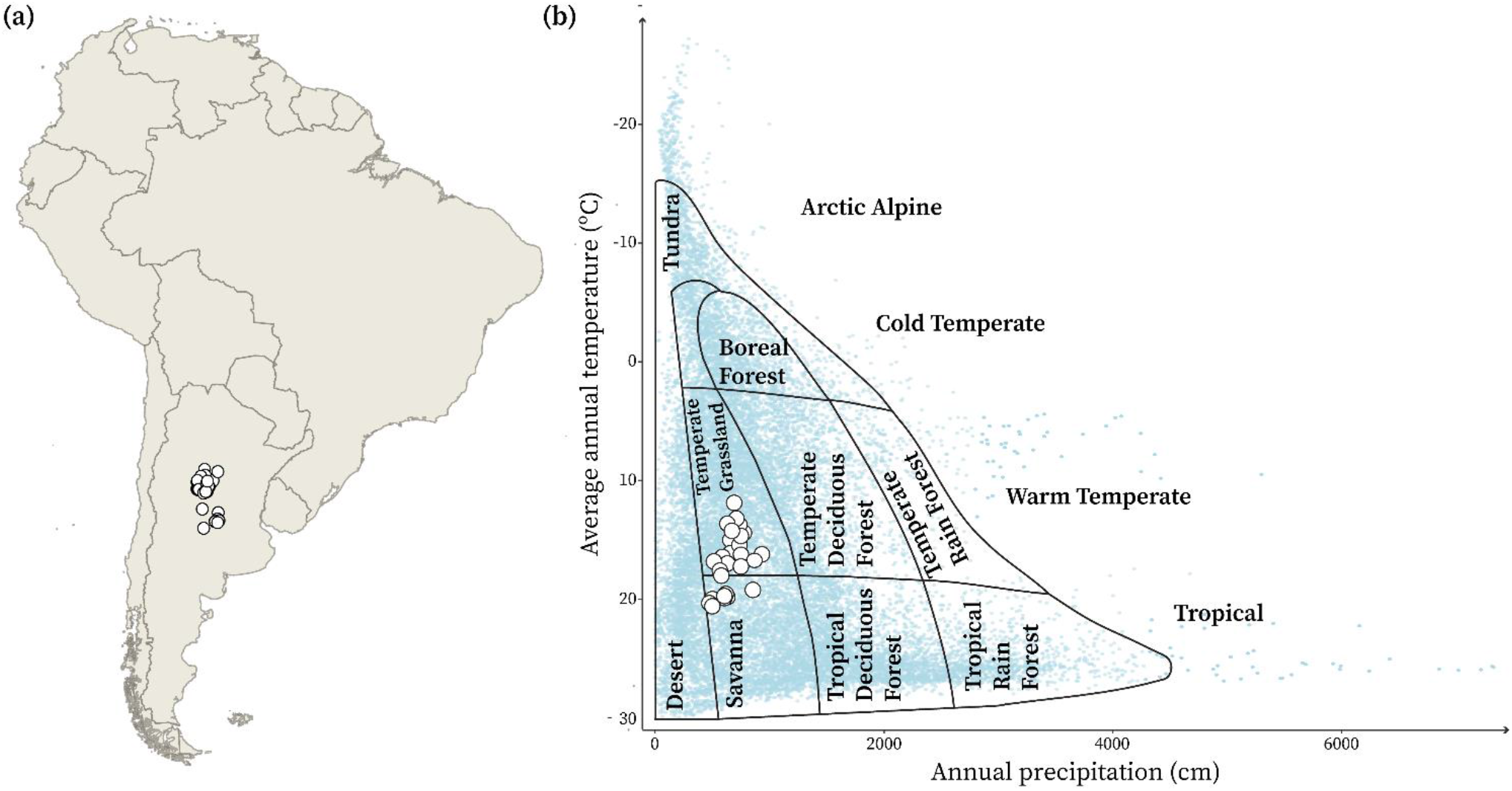
Geographical and climatic location of the plant communities studied (a) within the map of South America; and (b) in the Whittaker biome scheme (Whittaker, 1962). In (b), the white dots represent the Central Argentina communities, and the light-blue dots represent the locations of the species with information for the six traits used in the GSPFF (Díaz et al., 2016). The plant communities studied are located between the longitudes 29°37’32.6’ S and 34°52’2.9’ S and the latitudes 63°14’19.5’ W and 65°37’1’ W. The 40 points displayed in the panels are representative coordinates reflecting the diversity of precipitation and temperature across the 18 community types considered; see Table S1 for coordinates.

In this article, we set out to analyse the essential, ‘bare-bones’ functional trait space (hereafter phenospace, synonym of phenotypic space sensu Pigliucci, 2007) occupied by the vascular flora of Central Argentina, and to find out to what degree it matches the global phenospace represented by the GSPFF. We aimed to answer two main questions: First, do the regional and the global floras exhibit similar main axes of variation and trade-offs among traits? Our hypothesis is that these patterns are shared across both levels, as they are thought to be common across the majority of vascular plants (Díaz et al., 2016; Wright et al., 2004). Second, does the regional plant phenospace of Central Argentina support the habitat filtering hypothesis? According to this hypothesis, floras developed in relatively harsh climates should exhibit a smaller, as well as skewed phenospace than the one potentially allowed by the pool of species whose propagules are able to reach the site (Díaz et al., 1998a; Grime, 2006; Keddy, 1992; Swenson et al., 2012; Woodward & Diament, 1991). Central Argentina provides an ideal setting for examining functional diversity response to habitat filtering: Its climate and soils include some moderate to harsh conditions with clear habitat filtering potential (Fig. 1b). At the same time, it presents minimal species-pool limitations due to its lack of insularity that could seriously limit dispersion (Fig. 1a; Díaz et al., 1998), and its position at the interface of different phytogeographical regions (see below). Under the habitat filtering hypothesis, therefore, we expected the phenospace occupied by this regional flora to be significantly smaller (less functionally diverse) and more skewed towards conservative syndromes than the GSPFF. Moreover, it should be smaller and more skewed than expected by random on the basis of its species richness. More generally, through this analysis, we aimed to contribute to a better understanding of the factors that shape functional diversity at sub-global scales.

## 2 MATERIALS AND METHODS

### 2.1 Study area

The study area is located in Córdoba Province, Central Argentina (Fig. 1). To select a list of species representative of its flora, we used 793 highly detailed preexisting vegetation surveys carried out across 18 different plant community types within the Central Argentina floristic region (Cabido et al., 1994, 2018; Cantero et al., 2001; Giorgis et al., 2017), partially representing the Chaco, Espinal, and Pampeana Phytogeographic Provinces *sensu* Cabrera (Cabrera, 1953, see Table S1 for the full list of community types). In addition to being a phytogeographical ‘meeting point’, the region contains a wide range of ecosystems, encompassing forests, shrublands, and grasslands (Burkart et al., 1999; Cabido et al., 2018), with an altitudinal gradient spanning from c. 100 to c. 2800 m above sea level, and involves an old mountain system and lowlands of Pleistocene and Holocene sediments (Carignano & Cioccale, 2014). The mean annual precipitation ranges from 450 to 950 mm, and the mean temperature is between 10°C and 20°C (Fick & Hijmans, 2017). Soil fertility varies substantially across habitats, from Pampeana grasslands and Espinal forests, with some of the most fertile loess-based soils in the world, to the moderately fertile soils of Chaco forests, to seasonally flooded soils, to the extremely low fertility of highly rocky and sandy soils of some mountain grasslands, and the saline soils of the salt flats. For a better representation of the full richness of the native flora, we excluded severely disturbed communities, including only those with low to moderate known human use since the mid-1950s.

### 2.2 Species selection

From a total of 1297 vascular plant species recorded in 793 pre-existing field surveys across 18 different plant community types (Cabido et al., 1994, 2018; Cantero et al., 2001; Giorgis et al., 2017), we selected the most abundant ones in each community (highest percentage of ground cover) under the assumption that those are the ones that better reflect the prevailing environmental filters (Cingolani et al., 2007; Díaz et al., 1998a; Grime, 2006) and exert the largest influence on ecosystem properties (Díaz et al., 2007; Garnier et al., 2004; Pérez-Harguindeguy et al., 2013). The selection was conducted as follows: we averaged all available surveys for each community type and ranked species from highest to lowest relative cover. We then selected all species necessary to reach at least 60% of the total relative cover, starting with the most dominant. In addition, we included species that are common in the region and belong to groups underrepresented in the GSPFF (Díaz et al., 2016), such as ferns, palms, bromeliads, and succulents. Although they are in general of small size and tend not to reach high abundance except in specific localities, they are frequent and highly distinctive elements of the regional flora (Cabido et al., 2018; Cantero et al., 2001). Exotic species present in some of the sites were excluded from analysis. This selection process resulted in 410 priority species (see Table S2 for full list).

### 2.3 Trait measurements

We focused on the six functional traits used in the GSPFF (Díaz et al., 2016), which are known to be at the core of growth, survival, and reproduction of the vast majority of vascular plants and are at the same time available or relatively easy to measure for a large number of species worldwide (Pérez-Harguindeguy et al., 2013). These traits were: adult plant height (H, units: m), stem specific density (dry mass per unit of fresh stem volume, SSD, units: mg.mm^-3^), leaf area (one-sided surface area of an individual lamina or leaflet, LA, units: mm^2^), leaf mass per area (leaf dry mass per unit of lamina surface area, LMA, units: g.m^-2^), leaf nitrogen content per unit of lamina dry mass (Nmass, units: mg.g^-1^) and diaspore mass (mass of an individual seed or spore, SM, units: mg). When available, pre-existing data in the CORDOBASE database were used (Díaz et al., 2004 and unpublished data). The missing traits were measured afresh in the field for this study, following the methods by Pérez-Harguindeguy et al. (2013). The final dataset comprised 2045 cells of information on trait means (each the product of several replicates), of which 556 cells (27%) correspond to new primary data from the field. No trait information was imputed.

### 2.4 Data analysis and projection

We compared trait means between the regional flora of Central Argentina (hereafter ‘regional flora’), represented by our 410 selected species, and the global flora represented by the dataset published by Díaz et al. (2022) which corresponds to the same dataset used for the construction of the GSPFF (Díaz et al., 2016), and is to date the largest and most accurate compilation of empirically observed vascular plant species mean traits. To avoid disproportionately representing Central Argentina within the global context and to use GSPFF as an external reference, we chose not to incorporate our own dataset into the GSPFF data.

We tested for significant differences between the regional and global floras in the log10-transformed traits, distinguishing between woody and non-woody species, using a phylogenetically informed ANOVA, via the ‘phytools’ and ‘PhyloMaker’ packages in R (Jin & Qian, 2019; Revell, 2012). We used a jitter and a dot plot with error bars for visualization (R Core Team, 2020). In order to characterize the relationships between pairs of traits in the regional and global floras, we performed Standardized Major Axes regressions (Warton et al., 2006) using the ‘smatr’ package in R (R Core Team, 2020; Warton et al., 2006). Additionally, we conducted a Multivariate Analysis of Variance (MANOVA) to test whether the vectors of trait means of the regional and global flora differed significantly, using the ‘manova’ function (R Core Team, 2020; Warton et al., 2006). To examine the concordance of the trait covariance matrices of the global and Central Argentina floras, we conducted a random skewers analysis (Cheverud & Marroig, 2007), using the ‘evolqg’ package in R (Melo et al., 2015). To characterize the phenospace occupied by the regional flora, we carried out a Principal Component Analysis (PCA) on a 268 species x 6 trait matrix (total number of species in the regional flora for which we had empirical measurements for all six traits of interest), using the R-function ‘princomp’ (R Core Team, 2020). All traits were log10-transformed before analysis.

In addition, in order to formally compare the phenospaces of the regional and global floras, the 268 regional species were projected onto the GSPFF using the PhenoSpace application (Segrestin et al., 2021), i.e., the GSPFF was not modified, instead it was used as a canvas, with the regional species placed over it. Subsequently, to test whether the observed regional phenospace was significantly less spread over the GSPFF and/or functionally skewed than expected by chance for its species richness, as compared to the observed global phenospace, we carried out two different analyses. First, we computed the convex hulls of the regional and global floras (=observed convex hulls) following Cornwell (2006), using the ‘chull’ function in R, designed to identify the vertices of the convex hull (R Core Team, 2020). Random floras were generated to assess whether the regional convex hull matched one expected by chance for the same number of species. All the species of the GSPFF (global flora) were used as the pool from which 1000 floras with 268 species were sampled, and their convex hulls calculated. We then used the ‘sf’ package of R (R Core Team, 2020), to calculate the percentage of coverage over the observed global convex hull achieved by each of the random convex hulls and also by the observed regional convex hull. To assess significance, we calculated the extreme quantiles (5th and 95th percentiles) of the distribution of random coverage percentages, comparing them to the observed percentage of coverage to determine if it differed significantly from what would be expected by chance. Secondly, we overlaid a grid on the GSPFF, and calculated the percentage of random convex hulls that contained each cell. On this basis, we determined the areas of the GSPFF expected to be covered by chance and compared them with the observed regional convex hull.

Taking as primary dataset the positions of the 268 Central Argentina species placed over the GSPFF of Díaz et al. (2016) calculated according to Segrestin et al. (2021), we performed a Kernel density estimation. The resulting heat map indicates the regional species occurrence probability within the broader context of the GSPFF. The ‘stat_density2d’ function in the ggplot2 package of R was employed for this purpose (R Core Team, 2020). Graphs were made using both a color map and contour lines, following Díaz et al. (2016). Additionally, to assess whether the position of the regional centroid deviated from what would be expected by chance, we calculated the centroid positions of 1000 random floras projected over the GSPFF, using the procedure for generating random floras described above. We then carried out an inclusion test to determine whether the regional centroid fell within the area occupied by the centroids of these random floras. This test was performed using the ‘hypervolume’ package in R, following the methodology described by Blonder et al. (2014).

Lastly, after projecting the regional flora onto the GSPFF, we defined the regional species that were functionally distinct at the global scale as the ones surpassing the 0.99 density probability isoline of the GSPFF. We also identified the functionally distinct species following the definition and method of Violle et al. (2017). For this purpose, we constructed a new general pool of species, comprising the regional flora plus the global flora. We calculated each species’ position in the GSPFF and computed the distance matrix. We calculated the mean distance from each species to all others and standardized by dividing it by the maximum pairwise distance recorded in the distance matrix. Species with a mean distance that exceeded the 99th quartile were classified as functionally distinct.

## 3 RESULTS

### 3.1 Functional attributes of the regional flora of Central Argentina

The range and mean values of the six traits measured on the 410 species of the regional flora included in our analysis showed several differences with respect to those of the global flora (Table 1 and Fig. 2). Central Argentina species were, on average, significantly shorter and smaller-leafed than those in the global flora. Woody species in the regional flora had higher Nmass, and non-woody ones had higher SSD and LMA than those in the global flora. Additionally, the highest values of SM, LA, and H recorded for regional species were markedly lower than those recorded for the global flora. In turn, the relationships between pairs of individual traits showed remarkable similarity with those in the global flora, with no difference in slope direction in 11 out of a total of 15 bivariate tests (Table S3).

**TABLE 1.**
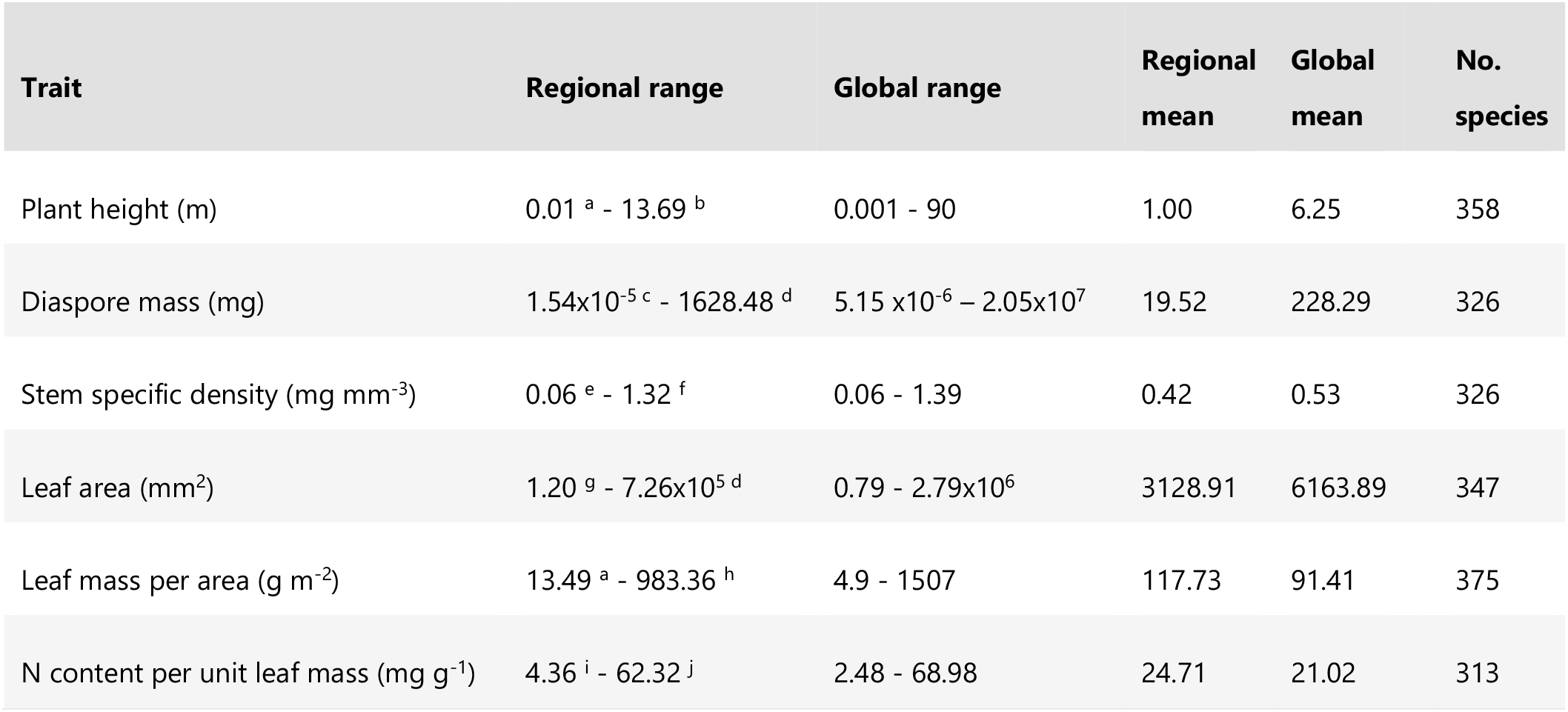
Distribution of key functional trait values of the regional flora of Central Argentina as compared with the global flora. Regional and global range and mean values for six functional traits, and the number of species in which they were measured in the regional flora. The information about global species was taken from Díaz et al. (2016) and Díaz et al. (2022). Species with minimum and maximum values for each trait in the regional flora are: a, *Salvinia nutans*; b, *Schinopsis lorentzii*; c, *Polystichum montevidense*; d, *Trithrinax campestris*; e, *Cleistocactus baumannii*; f, *Bulnesia foliosa*; g, *Strombocarpa strombulifera*; h, *Neltuma kuntzei*; i, *Tillandsia myosura*; j, *Datura ferox*.

**FIGURE 2.**
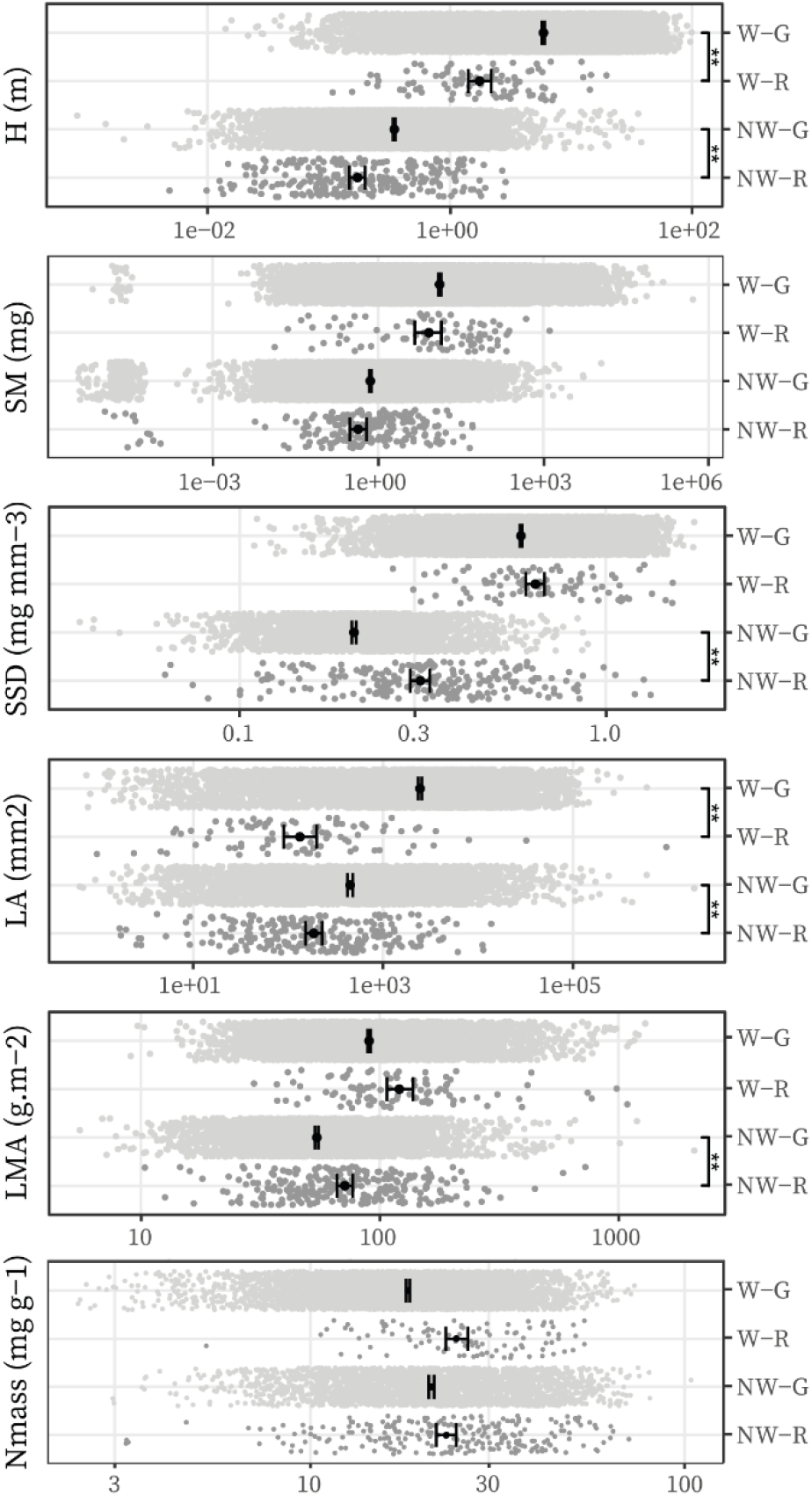
Distributions of the six traits in the regional flora of Central Argentina and the global flora represented in the GSPFF of Díaz et al. (2016), divided into woody and non-woody species. Each panel corresponds to one log-10 transformed trait, and includes a jitter plot, mean value, and standard deviation for each group. Asterisks indicate significant phylogenetically-informed differences in pairwise comparisons within the woody (W) and non-woody (NW) categories: **: p < 0.01, *: p< 0.05. W-G: Woody global species. W-R: Woody regional species. NW-G: Non-woody global species. NW-R: Non-woody regional species.

Of the 410 measured species of the regional flora, 268 had empirical information for all six traits considered. This group represents 20.66% of the total number of species reported for the study area (1297, Cabido et al., 1994, 2018; Cantero et al., 2001; Giorgis et al., 2017). Since the main criterion for species selection was abundance and/or frequency, the degree of representation of different taxonomic orders in our dataset varies, but we sampled >15% of each of the four taxonomic orders with >100 species according to the original surveys (Fig. S1, Cabido et al., 1994, 2018; Cantero et al., 2001; Giorgis et al., 2017). Species with complete and incomplete measurements for the six traits show similar distribution patterns for each trait (Fig. S2).

The MANOVA analysis showed that the vectors of trait means for the regional and global flora were significantly different (p-value < 0.001, Wilks’ lambda = 0.8). However, regarding whether trade-offs among traits in the regional flora were similar to those in the global one, the random skewers analysis showed a significant correlation between the trait covariance matrices of both floras, with a correlation coefficient of 0.93 and a standard deviation of 0.09. This result indicates that trait relationships in Central Argentina closely align with those observed globally. In line with this result, we found some strong similarities in main axes of variation between the two floras, as shown by the comparison between the PCA of Central Argentina species and that of global species (Table S4). As in the GSPFF, in the regional PCA plant stature and organ size primarily loaded onto the first principal component, while traits related to leaf resource economics predominantly loaded onto the second, with only slight differences in the raking of some traits. In particular, among the traits that loaded most heavily onto PC2, Nmass ranked higher than LA in our study.

### 3.2 The flora of Central Argentina against the backdrop of the GSPFF

Regarding whether the regional phenospace of Central Argentina supports the habitat filtering hypothesis by being *smaller* than the global flora, when projected onto the GSPFF, the 268 regional species showed a remarkable spread (Fig. 3a). The observed regional convex hull covered 68.75% of the observed global convex hull, not showing significant deviation from the *% coverage* expected by chance (Fig. 3b).

**FIGURE 3.**
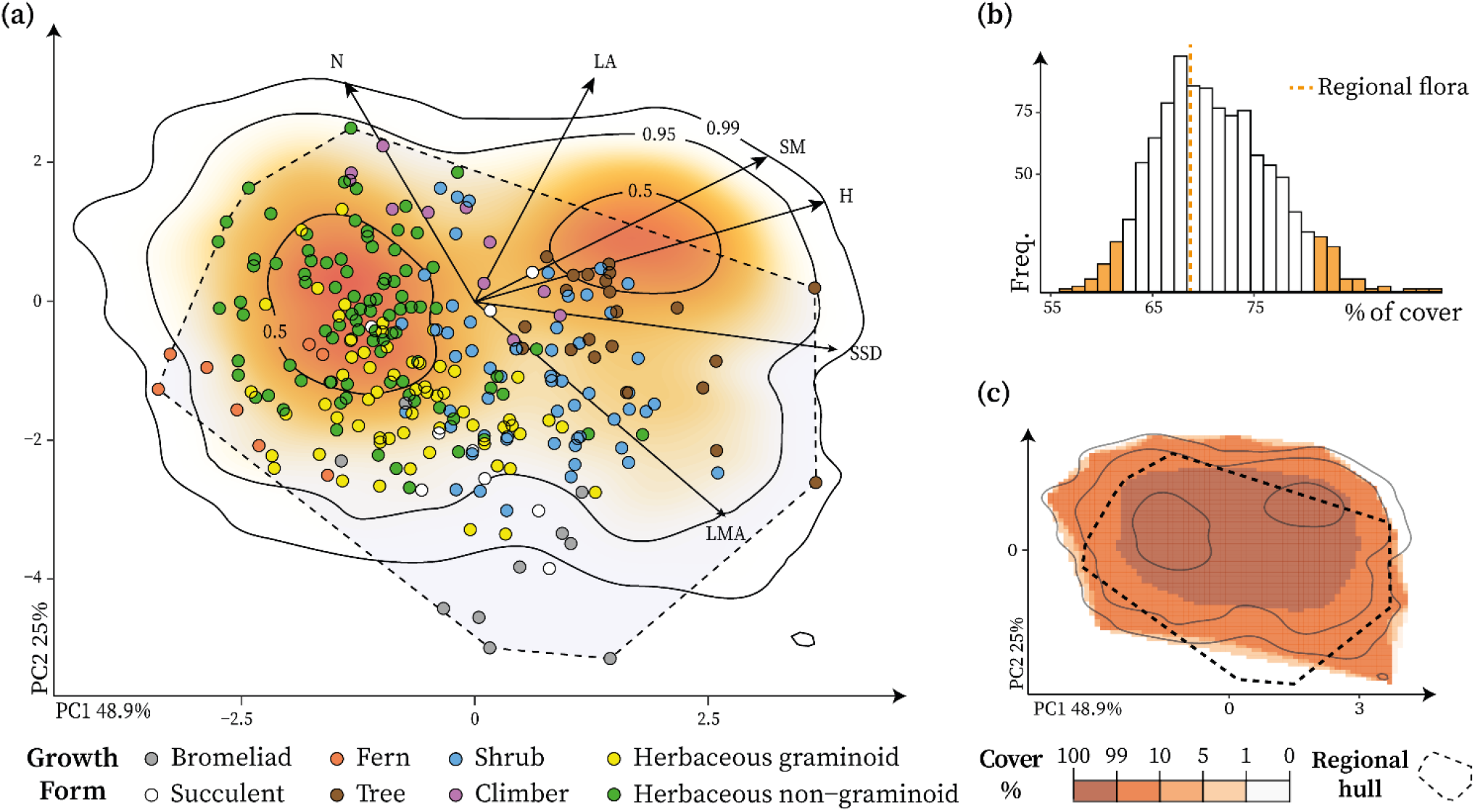
The regional flora of Central Argentina as compared with the global flora. (a) Regional species projected onto the GSPFF (Díaz et al., 2016): the panel displays the GSPFF as a heat map in the background; the GSPFF shows the distribution of 2214 species on the plane defined by principal component axes (PCA) 1 and 2, with isolines and colours representing probability density, ranging from high (dark orange) to low (light orange-white). The circular symbols overlaid indicate the positions of the 268 regional species with full trait information, color-coded by growth form; the black dashed line delineates the observed regional convex hull. (b) Convex hull coverage histogram: displaying the distribution of the percentage of overlap between the observed global convex hull and 1000 null regional convex hulls. The orange dashed line represents the percentage of overlap of the observed regional and global convex hulls. Orange bins denote the extreme quantiles (5th and 95th percentiles) of the random % coverage distribution. (c) Convex hull cover probability: the heat map represents the frequency in which the 1000 null convex hulls covered each cell of the grid in which we divided the GSPFF. The legend below the panel indicates the cover probability range from maximal (dark) to minimal (light). The observed regional convex hull (dashed line) is projected on top of the heat map.

However, regarding whether the regional phenospace of Central Argentina supports the habitat filtering hypothesis by being *skewed* relative to the global flora, the convex hull of the regional flora was significantly displaced with respect to random expectations (Fig. 3c). On the one hand, the upper-right corner, corresponding to tall trees with large-to moderately-sized leaves and seeds, densely populated in the GSPFF to the point of forming a functional hotspot (Díaz et al., 2016), appeared untapped by the regional flora. On the other hand, the Central Argentina flora extended towards conservativeness to an area beyond the GSPFF 0.99 isoline, which should have been empty.

In summary, the observed regional phenospace was more functionally skewed than expected by chance for its species number, but it was not significantly less spread over the GSPFF, because the *distribution* of the regional convex hull within the GSPFF deviated from expected, but its *% coverage* still aligned with expectations. This is because the reduction in *% coverage* due to the absence of tall trees is counterbalanced by an expansion of functional space due to the presence of extremely conservative species.

### 3.3 Functional ‘topography’ and globally distinct species in the flora of Central Argentina

Concerning whether the *distribution* of regional species over the GSPFF matches with the habitat filtering hypothesis, when projected onto the GSPFF, central Argentina species show notable differences in the *density of occupation* across different sectors of the global convex hull. By far, the most frequent species were non-woody with small leaves and seeds (Fig. 4a). These represent the only detectable functional hotspot in this regional flora, unlike the case of the GSPFF, which exhibits two prominent hotspots, one dominated by herbaceous plants and the other dominated by trees (indicated with dashed lines in the background to Fig. 4a, cfr. Díaz et al., 2016). The single hotspot of the Central Argentina flora partially overlapped with the herbaceous hotspot of the GSPFF, but appeared slightly displaced towards more conservative syndromes (higher LMA and lower Nmass, Fig. 4a). This displacement is because xerophytic graminoids and bromeliads are comparatively numerous and common in the regional flora. Although the ‘summit’ of species density (the hottest colour in the heat map of Fig. 4a) still overlapped with the herbaceous hotspot of the GSPFF and consisted almost entirely of herbaceous plants, the high-density area delimited by the 0.5 probability density isoline extended beyond it toward the woody sector of phenospace, and included a good number of shrubs and subshrubs (Fig. 4b). Besides, the regional flora centroid is different than expected by chance, since its position within the GSPFF falls outside the area occupied by the centroids of random floras according to the inclusion test performed (Fig. 4a). This indicates a distinct position of the regional flora, with its centroid shifted towards a smaller and more conservative syndrome.

**FIGURE 4.**
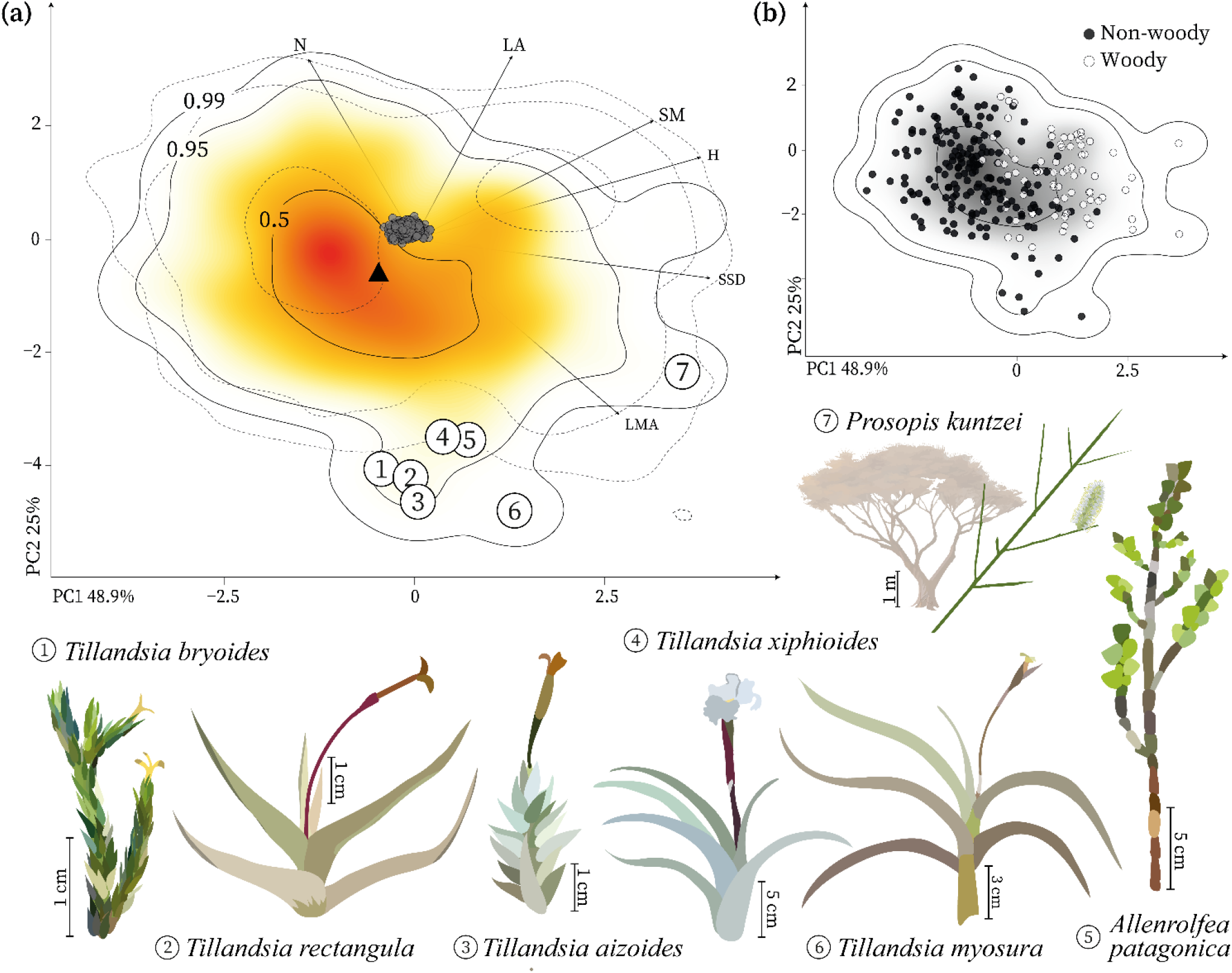
Common and distinct functional syndromes in the Central Argentina flora. (a) Kernel density estimation of the regional species: Projection of 268 vascular plant species representative of the regional flora onto the GSPFF (Díaz et al., 2016). The colour map indicates the kernel density estimation of the regional species, from highest (red) to lowest (white) occurrence probability. Contour lines indicate 0.5, 0.95, and 0.99 quantiles of species occurrence probability. Numbers in circles indicate the positions of the seven species identified as functionally distinct according to our and Violle et al.’s (2017) methodologies, illustrated around the main panel. In the inset background, the contour lines of 0.5, 0.95, and 0.99 quantiles of species occurrence probability of the GSPFF according to Díaz et al. (2016) are drawn with dotted lines. The black triangle marks the location of the regional centroid, and the cloud of black circles represent the centroid positions of 1000 floras expected by chance (samples of the global flora). (b) Woody and non-woody regional species distribution: distribution of woody (white circles) and non-woody (black circles) regional species depicted over the kernel density estimation of the regional species.

Regarding functional distinctiveness, six Central Argentina species fell outside the 0.99 density probability isoline of the GSPFF (species 1 to 6, Fig. 4a). Of these six regional species that we established as functionally distinct at the global scale, five are epiphytes of the *Tillandsia* genus (Bromeliaceae) and the remaining one, *Allenrolfea patagonica* (Amaranthaceae), grows only on highly saline soils. The application of Violle et al.’s (2017) methodology for the determination of globally distinct species yielded a similar result, except that it excluded *Allenrolfea patagonica* and the higher-statured *Tillandsia xiphioides* and added one more species, *Neltuma kuntzei* (Fabaceae), a xerophytic, thorny and leafless tree with photosynthetic stems (species 7, Fig. 4a).

## 4 DISCUSSION

### 4.1 The regional phenospace is consistent with the GSPFF

The phenospace occupied by the flora of Central Argentina shows a strong consistency in major axes, functional syndromes, and trait trade-offs with the GSPFF constructed by Díaz et al. (2016) based on a worldwide representative flora. The size of whole plant and plant organs on the one hand, and leaf resource economics on the other, and the three major traits determining PC1 and PC2 were identical in both floras. There were slight differences in the ranking of some of the traits in determining the PC axes due to the characteristics of this regional flora. In particular, among the traits more strongly determining PC2, Nmass ranked higher than LA in our study, which is consistent with the strong presence in the phytogeographical region of graminoids and bromeliads (low to extremely low Nmass) together with legumes (high Nmass except for the species with photosynthetic stems). In addition, the regional flora showed a much smaller range of LA, skewed towards smaller leaves, as could be expected in a temperate, semiarid flora. The predominance of legumes could also partially explain this pattern since the vast majority of the representatives of that family in the region have compound leaves, often with numerous very small leaflets.

The random skewers analysis showed a significant correlation between the trait covariance matrices of global and Central Argentina floras, and the correlation slopes between pairs of traits were remarkably similar. Of a total of 15 traits correlations, only four showed differences in slope direction between the two floras. All these differences can be explained by the predominance of moderately-to tall-statured woody legumes, which typically combine the capacity of symbiotically fixing N, compound deciduous leaves (thus low-LA, low-LMA, high-Nmass), and some of the largest seeds and densest woods in the region.

The consistency of the main axes of variation and trait trade-offs between the regional and global flora is compatible with our first hypothesis. Together with findings by Barajas-Barbosa et al. (2023), Thomas et al. (2020), and Carmona et al. (2021), among others, this result lends support to the robustness and universality of these fundamental functional dimensions across regions with different biogeographic history, climate, and insularity. The generality of the relationships between traits suggests that, while plant life displays a remarkable functional variability, it does so within constrained portions of the total trait space, tightly restrained by physiological limitations and interaction pressures (Wright et al., 2004). One major axis reflects the general pattern that the whole-plant size is coupled with the size of its organs, while the second axis reflects how the costs of leaf construction are typically coupled with its growth potential (Díaz et al., 2016). In addition, this consistency reinforces the GSPFF’s value as a useful tool for mapping plant species within a global context, enabling species comparisons and functional diversity analysis across diverse ecosystems.

### 4.2 The regional phenospace is remarkably wide and yet distinctive

The phenospace of the Central Argentina flora is remarkably wide, with almost 70% overlap with the GSPFF, despite containing almost ten times fewer species and having a much smaller climatic and geographic scope. This high regional dispersion contradicts, in principle, our second hypothesis, which predicted a significantly smaller phenospace due to habitat filtering. However, it coincides with Barajas-Barbosa et al. (2023), who found that the flora of the oceanic island of Tenerife, despite its high isolation, also showed an unexpectedly high functional richness.

Westoby et al. (2002) have pointed out that the spread of trait values within a local site is often as wide as that found over long regional gradients. Our findings, together with those of Barajas-Barbosa, lend support to these ideas and suggest that the pattern could be extended to broader geographical scales: the region of our study is much larger and more heterogeneous than a single site or even a single community type, but it still contains much less physical variation than the whole inhabited land area of the planet that is the scope of the GSPFF. Westoby et al. (2002) mentioned three potential explanations for the pattern they described: (a) strong microsite heterogeneity in physical conditions, (b) sustained immigration from source sites with different physical conditions into a sink site, and (c) game-theoretic or frequency-dependent processes. These are not mutually exclusive mechanisms and are quite difficult to disentangle in the field. A conclusion about their relative importance is well beyond the scope of our data and methods. However, we speculate that sustained immigration from source areas may not be the primary factor explaining the wide regional phenospace. While it cannot be entirely disregarded due to the region’s lack of insularity, we can question its importance since the floras of oceanic islands also show a remarkably high functional diversity, as described by Barajas-Barbosa et al. (2023) for Tenerife, and Díaz et al. (2007) for the mountain flora of the Hawaiian archipelago. Both studies suggest that the isolation that makes island floras relatively species-poor and taxonomically imbalanced does not pose equally severe restrictions on functional diversity. Therefore, while unrestricted immigration of propagules from source areas may contribute to the ample functional space of continental, well-connected areas like Central Argentina, isolation by itself does not appear to be a strong limitation for developing rich phenospaces in isolated areas.

Despite its great amplitude, the coverage of the global phenospace by the regional flora from Central Argentina is functionally skewed: tall trees with moderately sized to large leaves and seeds, which form a major hotspot in the GSPFF, are virtually absent. This absence is not an artefact of sampling, since we covered all the tall native tree species documented for the region; furthermore it is fully consistent with the climatic conditions, in particular the evapotranspiration/precipitation balance during the growing season and the occurrence of dry, relatively cold winters with frosts (Moles et al., 2009, 2014; Wright et al., 2017).

### 4.3 The regional phenospace has a shifted ‘topography’

The displacement of the regional centroid and functional hotspot towards more conservative syndromes with respect to the herbaceous hotspot of the GSPFF, its inclusion of a significant number of short shrub and subshrub growth forms, and its ‘functional encroachment’ into what is a relatively unpopulated sector in the GSPFF (Díaz et al., 2016), are also consistent with the climatic (seasonal or year-round drought) and edaphic (rocky, highly sandy, saline soils) conditions that prevail in some areas of Central Argentina. Xerophytic graminoids, bromeliads, shrubs, and subshrubs are also present in the GSPFF but are rarer than other growth forms.

A ‘shifted topography’ –coldspots or ‘valleys’ and hotspots or ‘peaks’ of recurrent combinations of trait values found in positions in phenospace markedly different from the global distribution of ‘peaks’ and ‘valleys’ shown in the GSPFF– has also been reported by other authors studying floras developed under harsh climates. Barajas-Barbosa et al. (2023) found a single functional hotspot dominated by shrubby rather than herbaceous growth forms. Thomas et al. (2020) found that, under the extremely cold conditions of the tundra, species occupied a wide area of functional space in terms of resource use economics but a highly constrained one in terms of stature and organ size, also with a strong presence of shrubs and subshrubs.

In agreement with the displacement of the regional functional hotspot towards more conservative syndromes, the species of the regional flora identified as functionally distinct are extremely conservative. Regardless of whether we use the GSPFF’s 0.99 isoline determination or Violle et al.’s (2017) criteria, the resulting functionally distinct species are all highly specialised stress-tolerators, rather than representative of particularly unique local lineages. *Allenrolfea patagonica* is a xero-halophyte succulent characteristic of Central and Northern Argentina’s saltflats (Pérez Cuadra & Hermann, 2014; POWO, 2023) and belongs to Amaranthaceae, a geographically extended family in which salt-tolerance is common. *Neltuma* is a widespread genus across the Americas with many representatives in Argentina’s flora (POWO, 2023; Zuloaga et al., 2008). However, the functionally distinct *Neltuma kuntzei* stands out as a particularly extreme case, even within the Central Argentina region, of adaptation to both drought and herbivory, with an extremely dense wood, most of its photosynthesis through the year being carried out by tough, thorn-like photosynthetic stems, and still maintaining the tree growth form. Finally, *Tillandsia* is a genus endemic to the whole of the Americas (POWO, 2023), whose members are all epiphytic or saxicolous rosettes with different degrees of xerophytism. The five *Tillandsia* species detected in this study as functionally distinct are markedly xerophytic even within the genus, with extreme adaptions to drought and nutrient scarcity, as manifested in their very high LMA and extremely low Nmass, as well as CAM photosynthetic metabolism (Benzing & Bennett, 2000; Crayn et al., 2015) and leaves densely covered by trichomes, which multiply the surface intercepting moisture and particles from the air (Estrella-Parra et al., 2019; Stefano et al., 2006).

## 5 CONCLUDING REMARKS

Summing up, in response to our first initial question, the regional flora of Central Argentina shows the same general axes of variation and trade-offs among traits as the global flora. This, together with recent works on regional floras developed under moderate to harsh environmental conditions in other areas of the world, lend support to the universality, among vascular plants, of the two major axes of functional variation represented by size and leaf resource economics.

In response to our second initial question, the shifted species density distribution, the ‘missing’ regions in the regional phenospace with respect to the global one, as well as the highly conservative syndromes of the functionally distinct species in the regional flora, are all consistent with the regional climate and soils. This consistency strongly suggests the operation of habitat filtering and thus our hypothesis appears to find partial support: although the prevailing environmental conditions do not appear strong enough to result in a significantly smaller regional phenospace, they are likely to be the reason behind the unequal filling and the displacement of functional hotspots in the regional phenospace, as compared to the global phenospace.

More generally, our findings and those of other studies explicitly using the GSPFF as a baseline allow for some speculation about the role and effects of habitat filtering. While more formal and geographically comprehensive testing would be in order, evidence from regions with contrasting climates and degrees of isolation suggests that, at least at the regional scale, habitat filtering shapes functional ‘topographies’ more than it constrains the extent of phenospaces. This allows for a rich range of syndromes to survive, but with some of them being much more common than others depending on the prevailing environmental conditions.

## Acknowledgements

We thank G. Arias, G. Bertone, N. Gallardo, and I. Lynch-Ianniello for technical support; D. Bianchi, M. L. Bruno, F. Fernández-Catinot, J. Medrano-Santos, and A. Sánchez for their assistance in the field; and to the Parque Provincial y Reserva Forestal Natural Chancaní and local landowners and landwellers for access to experimental field sites established in their land; to M. Cabido, J. J. Cantero, M. Giorgis, and R. Morero for extremely valuable taxonomic and overall advice and original vegetation surveys, and to Jens Kattge from TRY for data facilitation. We acknowledge the contribution of Helge Bruelheide and Tim Walther at the Geobotany lab at Martin-Luther-University Halle-Wittenberg, Germany, for leaf carbon and nitrogen content analyses. We are also thankful to M. L. Bruno and M. L. Lipoma for access to unpublished data, and to L. D. Gorné for insightful analytical advice. This work was supported by the ARBOLES project funded by NERC (NE/S011811/1), and the Red Federal de Alto Impacto CONATURAR, Argentina (2023-102072649-APN-MCT).

## Authors contributions

Both authors together conceived and designed methodology, collected and analysed the data, and wrote the manuscript.

