## Supplementary material for "Functional diversity of a temperate continental flora: Wide but skewed coverage of the global spectrum of plant form and function"

### SUPPORTING INFORMATION

**TABLE S1** Table of communities studied. The table presents the 18 community types analyzed, along with their corresponding Phytogeographical Province affiliation (following Cabrera, 1953) and representative coordinates. For community types with a wide distribution, multiple coordinates are provided.

| Community type | Phytogeographical province | Representative coordinates | |
|---|---|---|---|
| | | Latitude | Longitude |
| Grassland | Pampeana | 33º28′17.2′′S | 63º19′49.2′′W |
| | | 34º02′25.1′′S | 63º23′38.6′′W |
| Sub-halophile shrubland | Pampeana | 34º17′43.5′′S | 63º25′10.3′′W |
| Halophile shrubland | Pampeana | 34º20′14.3′′S | 63º23′47.4′′W |
| Seasonally-flooded communities dominated by *Stipa densiflora* | Pampeana | 34º11′02.2′′S | 63º14′19.5′′W |
| Seasonally-flooded communities of reeds | Pampeana | 34º15′57.0′′S | 63º39′34.6′′W |
| Freshwater spring | Pampeana | 34º16′35.8′′S | 63º38′10.5′′W |
| Halophile grassland | Pampeana | 34º21′36.1′′S | 63º23′01.0′′W |
| Forest dominated by *Aspidosperma quebracho-blanco* and *Neltuma flexuosa* | Chaco | 30º41′09.4′′S | 64º42′18.8′′W |
| | | 30º42′41.3′′S | 64º44′20.6′′W |
| | | 30º49′54.4′′S | 64º57′08.7′′W |
| | | 31º22′28.4′′S | 65º27′15.1′′W |
| | | 31º20′31.7′′S | 65º35′11.0′′W |
| | | 31º23′53.2′′S | 65º35′19.5′′W |
| Shrubland dominated by *Larrea divaricata* and *Mimozyganthus carinatus* | Chaco | 31º21′46.8′′S | 65º37′01.0′′W |
| | | 31º18′52.1′′S | 65º34′34.9′′W |
| Open shrubland dominated by *Larrea divaricata* | Chaco | 30º48′40.6′′S | 65º26′10.5′′W |
| | | 31º26′32.8′′S | 65º28′56.4′′W |
| Forest and shrubland with *Stetsonia coryne* | Chaco | 30º44′36.0′′S | 65º36′12.4′′W |
| | | 29º37′32.6′′S | 64º48′55.9′′W |
| Ecotone between forest and saltflats | Chaco | 30º09′04.1′′S | 64º45′08.7′′W |
| | | 30º35′28.6′′S | 65º31′59.6′′W |
| Mountain forest | Chaco | 30º45′08.9′′S | 64º03′07.8′′W |
| | | 30º37′35.8′′S | 63º55′37.2′′W |
| Forest dominated by *Schinopsis balansae* | Chaco | 29º49′10.4′′S | 63º26′13.9′′W |
| Forest dominated by *Neltuma caldenia* | Espinal | 34º52′02.9′′S | 64º54′23.4′′W |
| Mountain shrubland | Chaco | 30º52′56.5′′S | 64º37′12.5′′W |
| | | 31º10′31.0′′S | 64º20′23.7′′W |
| | | 31º30′06.0′′S | 64º34′53.1′′W |
| Mountain grassland | Chaco | 31º36′11.0′′S | 64º32′42.3′′W |
| | | 33º09′35.4′′S | 65º03′25.1′′W |
| | | 31º08′13.3′′S | 64º24′55.2′′W |

| | | | |
|---|---|---|---|
| Rock outcrops | Pampeana, Chaco | 31°35′41.3′′S<br>31°36′18.5′′S<br>30°40′30.3′′S | 64°42′49.7′′W<br>64°40′20.2′′W<br>64°30′14.1′′W |

**TABLE S2** Table of the Central Argentina vascular plant species included in this study. The table contains all the species for which at least one trait was measured. In column Complete, 1 indicates the species for which all six traits of interest were measured and, therefore, were included in the PCA and convex hull analyses. The nomenclature follows Zuloaga et al. (2008).

| Species | Family | Growth form | Complete |
|---|---|---|---|
| *Abutilon grandifolium* | Malvaceae | Herbaceous non-graminoid | 1 |
| *Acalypha communis* | Euphorbiaceae | Herbaceous non-graminoid | 1 |
| *Acanthospermum hispidum* | Asteraceae | Herbaceous non-graminoid | 0 |
| *Acanthostyles buniifolius* | Asteraceae | Shrub | 1 |
| *Achyrocline satureioides* | Asteraceae | Shrub | 1 |
| *Acicarpha tribuloides* | Calyceraceae | Herbaceous non-graminoid | 1 |
| *Acmella decumbens* | Asteraceae | Herbaceous non-graminoid | 0 |
| *Acmella leptophylla* | Asteraceae | Herbaceous non-graminoid | 0 |
| *Adesmia bicolor* | Fabaceae | Herbaceous non-graminoid | 0 |
| *Adesmia corymbosa* | Fabaceae | Herbaceous non-graminoid | 0 |
| *Adiantopsis chlorophylla* | Papaveraceae | Fern | 0 |
| *Adiantum raddianum* | Pteridaceae | Fern | 1 |
| *Adiantum thalictroides* | Pteridaceae | Fern | 1 |
| *Agrostis montevidensis* | Poaceae | Herbaceous graminoid | 1 |
| *Alchemilla pinnata* | Rosaceae | Herbaceous non-graminoid | 1 |
| *Allenrolfea patagonica* | Amaranthaceae | Leaf succulent | 1 |
| *Allionia incarnata* | Nyctaginaceae | Herbaceous non-graminoid | 1 |
| *Aloysia gratissima* | Verbenaceae | Shrub | 1 |
| *Alternanthera albida* | Amaranthaceae | Herbaceous non-graminoid | 0 |
| *Alternanthera pumila* | Amaranthaceae | Herbaceous non-graminoid | 0 |
| *Alternanthera pungens* | Amaranthaceae | Herbaceous non-graminoid | 1 |
| *Amaranthus hybridus* | Amaranthaceae | Herbaceous non-graminoid | 1 |
| *Amauropelta argentina* | Thelypteridaceae | Fern | 0 |
| *Ambrosia tenuifolia* | Asteraceae | Herbaceous non-graminoid | 1 |
| *Amphilophium carolinae* | Bignoniaceae | Climber | 1 |
| *Anemia australis* | Anemiaceae | Fern | 1 |
| *Anemia tomentosa* | Anemiaceae | Fern | 0 |
| *Anredera cordifolia* | Basellaceae | Climber | 1 |
| *Araujia brachystephana* | Apocynaceae | Climber | 1 |

| | | | |
|---|---|---|---|
| *Argyrochosma nivea* | Pteridaceae | Fern | 1 |
| *Aristida achalensis* | Poaceae | Herbaceous graminoid | 1 |
| *Aristida adscensionis* | Poaceae | Herbaceous graminoid | 1 |
| *Aristida mendocina* | Poaceae | Herbaceous graminoid | 1 |
| *Aristida spegazzinii* | Poaceae | Herbaceous graminoid | 1 |
| *Aristolochia argentina* | Aristolochiaceae | Climber | 1 |
| *Asplenium gilliesii* | Aspleniaceae | Fern | 0 |
| *Asplenium resiliens* | Aspleniaceae | Fern | 1 |
| *Astragalus bergii* | Fabaceae | Herbaceous non-graminoid | 0 |
| *Astragalus parodii* | Fabaceae | Herbaceous non-graminoid | 0 |
| *Atamisquea emarginata* | Capparaceae | Shrub | 1 |
| *Atriplex argentina* | Amaranthaceae | Shrub | 1 |
| *Atriplex cordubensis* | Amaranthaceae | Herbaceous non-graminoid | 0 |
| *Atriplex lampa* | Amaranthaceae | Shrub | 0 |
| *Atriplex undulata* | Amaranthaceae | Shrub | 0 |
| *Austroblechnum penna-marina* | Blechnaceae | Fern | 0 |
| *Austroeupatorium inulifolium* | Asteraceae | Herbaceous non-graminoid | 0 |
| *Austroflourensia thurifera* | Asteraceae | Shrub | 1 |
| *Ayenia cordobensis* | Malvaceae | Herbaceous non-graminoid | 1 |
| *Baccharis aliena* | Asteraceae | Shrub | 1 |
| *Baccharis articulata* | Asteraceae | Shrub | 1 |
| *Baccharis coridifolia* | Asteraceae | Shrub | 0 |
| *Baccharis linearifolia* | Asteraceae | Shrub | 1 |
| *Baccharis tucumanensis* | Asteraceae | Shrub | 1 |
| *Baccharis ulicina* | Asteraceae | Shrub | 0 |
| *Berberis hieronymi* | Berberidaceae | Shrub | 1 |
| *Bidens andicola* | Asteraceae | Herbaceous non-graminoid | 1 |
| *Bidens pilosa* | Asteraceae | Herbaceous non-graminoid | 1 |
| *Bidens subalternans* | Asteraceae | Herbaceous non-graminoid | 1 |
| *Bidens triplinervia* | Asteraceae | Herbaceous non-graminoid | 0 |
| *Blechnum auriculatum* | Blechnaceae | Fern | 0 |
| *Blechnum laevigatum* | Blechnaceae | Fern | 0 |
| *Blumenbachia hieronymi* | Loasaceae | Herbaceous non-graminoid | 0 |
| *Boerhavia cordobensis* | Nyctaginaceae | Herbaceous non-graminoid | 1 |
| *Boerhavia diffusa* | Nyctaginaceae | Herbaceous non-graminoid | 1 |
| *Borreria verticillata* | Rubiaceae | Herbaceous non-graminoid | 1 |
| *Bothriochloa laguroides* | Poaceae | Herbaceous graminoid | 1 |
| *Bothriochloa saccharoides* | Poaceae | Herbaceous graminoid | 1 |
| *Botrychium australe* | Ophioglossaceae | Fern | 0 |
| *Bouteloua aristidoides* | Poaceae | Herbaceous graminoid | 1 |
| *Bouteloua curtipendula* | Poaceae | Herbaceous graminoid | 1 |
| *Bromus auleticus* | Poaceae | Herbaceous graminoid | 1 |
| *Bromus catharticus* | Poaceae | Herbaceous graminoid | 0 |
| *Buddleja stachyoides* | Scrophulariaceae | Shrub | 1 |

| | | | |
|---|---|---|---|
| *Bulbostylis funckii* | Cyperaceae | Herbaceous graminoid | 0 |
| *Bulbostylis juncoides* | Cyperaceae | Herbaceous graminoid | 0 |
| *Bulnesia foliosa* | Zygophyllaceae | Shrub | 1 |
| *Bulnesia retama* | Zygophyllaceae | Shrub | 1 |
| *Callianthe pauciflora* | Malvaceae | Herbaceous non-graminoid | 1 |
| *Cantinoa mutabilis* | Lamiaceae | Herbaceous non-graminoid | 1 |
| *Capsicum chacoënse* | Solanaceae | Herbaceous non-graminoid | 1 |
| *Cardionema ramosissimum* | Caryophyllaceae | Herbaceous non-graminoid | 1 |
| *Carex distenta* | Cyperaceae | Herbaceous graminoid | 1 |
| *Carex fuscula* | Cyperaceae | Herbaceous graminoid | 1 |
| *Carex phleoides* | Cyperaceae | Herbaceous graminoid | 0 |
| *Castela coccinea* | Simaroubaceae | Shrub | 1 |
| *Celtis ehrenbergiana* | Cannabaceae | Shrub | 1 |
| *Celtis tala* | Cannabaceae | Tree | 1 |
| *Cenchrus spinifex* | Poaceae | Herbaceous graminoid | 0 |
| *Cereus aethiops* | Cactaceae | Stem succulent | 1 |
| *Cereus forbesii* | Cactaceae | Stem succulent | 1 |
| *Cestrum parqui* | Solanaceae | Shrub | 1 |
| *Chaerophyllum andicola* | Apiaceae | Herbaceous non-graminoid | 1 |
| *Chaptalia integerrima* | Asteraceae | Herbaceous non-graminoid | 1 |
| *Chaptalia nutans* | Asteraceae | Herbaceous non-graminoid | 1 |
| *Chascolytrum subaristatum* | Poaceae | Herbaceous graminoid | 1 |
| *Cheilanthes buchtienii* | Pteridaceae | Fern | 1 |
| *Chevreulia sarmentosa* | Asteraceae | Herbaceous non-graminoid | 0 |
| *Chloris virgata* | Poaceae | Herbaceous graminoid | 1 |
| *Cinnagrostis hieronymi* | Poaceae | Herbaceous graminoid | 1 |
| *Cleistocactus baumannii* | Cactaceae | Stem succulent | 1 |
| *Clematis montevidensis* | Ranunculaceae | Climber | 1 |
| *Clinopodium gilliesii* | Lamiaceae | Shrub | 0 |
| *Colletia spinosissima* | Rhamnaceae | Shrub | 1 |
| *Cologania broussonetii* | Fabaceae | Herbaceous non-graminoid | 1 |
| *Commelina erecta* | Commelinaceae | Herbaceous non-graminoid | 1 |
| *Condalia buxifolia* | Rhamnaceae | Tree | 1 |
| *Condalia microphylla* | Rhamnaceae | Shrub | 1 |
| *Condalia montana* | Rhamnaceae | Shrub | 1 |
| *Conyza bonariensis* | Asteraceae | Herbaceous non-graminoid | 1 |
| *Cordobia argentea* | Malpighiaceae | Climber | 1 |
| *Cortaderia speciosa* | Poaceae | Herbaceous graminoid | 1 |
| *Cortesia cuneifolia* | Boraginaceae | Shrub | 1 |
| *Cotula mexicana* | Asteraceae | Herbaceous non-graminoid | 0 |
| *Crassula peduncularis* | Crassulaceae | Herbaceous non-graminoid | 0 |
| *Cressa nudicaulis* | Convolvulaceae | Herbaceous non-graminoid | 1 |
| *Croton argentinus* | Euphorbiaceae | Shrub | 1 |
| *Croton lachnostachyus* | Euphorbiaceae | Shrub | 1 |

| | | | |
|---|---|---|---|
| *Cucurbitella asperata* | Cucurbitaceae | Climber | 1 |
| *Cuphea glutinosa* | Lythraceae | Herbaceous non-graminoid | 0 |
| *Cyclolepis genistoides* | Asteraceae | Shrub | 1 |
| *Cynodon dactylon* | Poaceae | Herbaceous graminoid | 1 |
| *Cynophalla retusa* | Capparaceae | Tree | 0 |
| *Cyperus reflexus* | Cyperaceae | Herbaceous graminoid | 1 |
| *Datura ferox* | Solanaceae | Herbaceous non-graminoid | 1 |
| *Daucus pusillus* | Apiaceae | Herbaceous non-graminoid | 1 |
| *Deinacanthon urbanianum* | Bromeliaceae | Bromeliad | 1 |
| *Desmodium uncinatum* | Fabaceae | Climber | 0 |
| *Dichondra microcalyx* | Convolvulaceae | Herbaceous non-graminoid | 1 |
| *Dichondra repens* | Convolvulaceae | Herbaceous non-graminoid | 1 |
| *Dichondra sericea* | Convolvulaceae | Herbaceous non-graminoid | 1 |
| *Dicliptera squarrosa* | Acanthaceae | Herbaceous non-graminoid | 0 |
| *Digitaria californica* | Poaceae | Herbaceous graminoid | 1 |
| *Digitaria insularis* | Poaceae | Herbaceous graminoid | 0 |
| *Distichlis acerosa* | Poaceae | Herbaceous graminoid | 1 |
| *Distichlis spicata* | Poaceae | Herbaceous graminoid | 0 |
| *Dolichandra cynanchoides* | Bignoniaceae | Climber | 1 |
| *Doryopteris lorentzii* | Pteridaceae | Fern | 0 |
| *Dyckia floribunda* | Bromeliaceae | Bromeliad | 1 |
| *Echinopsis leucantha* | Cactaceae | Stem succulent | 0 |
| *Elaphoglossum gayanum* | Lomariopsidaceae | Fern | 1 |
| *Eleocharis pseudoalbibracteata* | Cyperaceae | Herbaceous graminoid | 1 |
| *Ephedra triandra* | Ephedraceae | Shrub | 1 |
| *Equisetum giganteum* | Equisetaceae | Herbaceous non-graminoid | 0 |
| *Eragrostis lugens* | Poaceae | Herbaceous graminoid | 1 |
| *Eragrostis retinens* | Poaceae | Herbaceous graminoid | 0 |
| *Eriosema edule* | Fabaceae | Herbaceous non-graminoid | 0 |
| *Eryngium agavifolium* | Apiaceae | Herbaceous non-graminoid | 1 |
| *Eryngium horridum* | Apiaceae | Herbaceous non-graminoid | 1 |
| *Eryngium nudicaule* | Apiaceae | Herbaceous non-graminoid | 1 |
| *Erythrostemon gilliesii* | Fabaceae | Shrub | 1 |
| *Euphorbia hyssopifolia* | Euphorbiaceae | Herbaceous non-graminoid | 0 |
| *Euphorbia serpens* | Euphorbiaceae | Herbaceous non-graminoid | 1 |
| *Eustachys retusa* | Poaceae | Herbaceous graminoid | 1 |
| *Evolvulus arizonicus* | Convolvulaceae | Herbaceous non-graminoid | 1 |
| *Evolvulus sericeus* | Convolvulaceae | Herbaceous non-graminoid | 1 |
| *Festuca dissitiflora* | Poaceae | Herbaceous graminoid | 0 |
| *Festuca hieronymi* | Poaceae | Herbaceous graminoid | 1 |
| *Festuca lilloi* | Poaceae | Herbaceous graminoid | 1 |
| *Flaveria bidentis* | Asteraceae | Herbaceous non-graminoid | 1 |
| *Fleischmannia prasiifolia* | Asteraceae | Herbaceous non-graminoid | 0 |
| *Galactia marginalis* | Fabaceae | Herbaceous non-graminoid | 0 |

| | | | |
|---|---|---|---|
| *Galium richardianum* | Rubiaceae | Herbaceous non-graminoid | 1 |
| *Gamochaeta filaginea* | Asteraceae | Herbaceous non-graminoid | 1 |
| *Gaultheria poeppigii* | Ericaceae | Shrub | 0 |
| *Gentianella multicaulis* | Gentianaceae | Herbaceous non-graminoid | 0 |
| *Gentianella parviflora* | Gentianaceae | Herbaceous non-graminoid | 1 |
| *Geoffroea decorticans* | Fabaceae | Tree | 1 |
| *Geranium magellanicum* | Geraniaceae | Herbaceous non-graminoid | 0 |
| *Geranium parodii* | Geraniaceae | Herbaceous non-graminoid | 0 |
| *Glandularia dissecta* | Verbenaceae | Herbaceous non-graminoid | 1 |
| *Glandularia peruviana* | Verbenaceae | Herbaceous non-graminoid | 1 |
| *Gomphrena martiana* | Amaranthaceae | Herbaceous non-graminoid | 0 |
| *Gomphrena pulchella* | Amaranthaceae | Herbaceous non-graminoid | 1 |
| *Gomphrena tomentosa* | Amaranthaceae | Herbaceous non-graminoid | 1 |
| *Gouinia paraguayensis* | Poaceae | Herbaceous graminoid | 1 |
| *Grahamia bracteata* | Portulacaceae | Shrub | 1 |
| *Grindelia globularifolia* | Asteraceae | Herbaceous non-graminoid | 0 |
| *Grindelia pulchella* | Asteraceae | Shrub | 1 |
| *Gymnocalycium monvillei* | Cactaceae | Stem succulent | 1 |
| *Harrisia pomanensis* | Cactaceae | Stem succulent | 0 |
| *Heterostachys ritteriana* | Chenopodiaceae | Shrub | 1 |
| *Hieracium gigantium* | Asteraceae | Herbaceous non-graminoid | 1 |
| *Hydrocotyle bonariensis* | Araliaceae | Herbaceous non-graminoid | 0 |
| *Hypochaeris argentina* | Asteraceae | Herbaceous non-graminoid | 1 |
| *Hypochaeris caespitosa* | Asteraceae | Herbaceous non-graminoid | 1 |
| *Hypoxis humilis* | Hypoxidaceae | Herbaceous graminoid | 1 |
| *Ipomoea purpurea* | Convolvulaceae | Climber | 1 |
| *Iresine diffusa* | Amaranthaceae | Herbaceous non-graminoid | 0 |
| *Jarava ichu* | Poaceae | Herbaceous graminoid | 1 |
| *Jarava juncoides* | Poaceae | Herbaceous graminoid | 1 |
| *Jarava pseudoichu* | Poaceae | Herbaceous graminoid | 1 |
| *Jodina rhombifolia* | Santalaceae | Tree | 1 |
| *Juncus balticus* | Juncaceae | Herbaceous graminoid | 1 |
| *Juncus microcephalus* | Juncaceae | Herbaceous graminoid | 1 |
| *Juncus pallescens* | Juncaceae | Herbaceous graminoid | 1 |
| *Juncus uruguensis* | Juncaceae | Herbaceous graminoid | 1 |
| *Justicia squarrosa* | Acanthaceae | Herbaceous non-graminoid | 1 |
| *Justicia tweediana* | Acanthaceae | Herbaceous non-graminoid | 0 |
| *Justicia xylosteoides* | Acanthaceae | Shrub | 1 |
| *Krapovickasia flavescens* | Malvaceae | Herbaceous non-graminoid | 1 |
| *Laennecia sophiifolia* | Asteraceae | Herbaceous non-graminoid | 0 |
| *Lantana camara* | Verbenaceae | Shrub | 1 |
| *Lantana grisebachii* | Verbenaceae | Herbaceous non-graminoid | 0 |
| *Larrea cuneifolia* | Zygophyllaceae | Shrub | 1 |
| *Larrea divaricata* | Zygophyllaceae | Shrub | 1 |

| | | | |
|---|---|---|---|
| *Lepechinia floribunda* | Lamiaceae | Shrub | 0 |
| *Lepidium bonariense* | Brassicaceae | Herbaceous non-graminoid | 0 |
| *Lepidium didymum* | Brassicaceae | Herbaceous non-graminoid | 0 |
| *Leptochloa crinita* | Poaceae | Herbaceous graminoid | 1 |
| *Leptochloa pluriflora* | Poaceae | Herbaceous graminoid | 1 |
| *Ligaria cuneifolia* | Loranthaceae | Herbaceous non-graminoid | 1 |
| *Lippia salsa* | Verbenaceae | Shrub | 1 |
| *Lippia turbinata* | Verbenaceae | Herbaceous non-graminoid | 0 |
| *Lithraea molleoides* | Anacardiaceae | Tree | 1 |
| *Lobelia hederacea* | Campanulaceae | Herbaceous non-graminoid | 0 |
| *Lorentzianthus viscidus* | Asteraceae | Shrub | 0 |
| *Lucilia acutifolia* | Asteraceae | Herbaceous non-graminoid | 0 |
| *Lycium chilense* | Solanaceae | Shrub | 1 |
| *Lycium elongatum* | Solanaceae | Shrub | 1 |
| *Lycium tenuispinosum* | Solanaceae | Shrub | 1 |
| *Lycopodium clavatum* | Lycopodiaceae | Herbaceous non-graminoid | 0 |
| *Maihueniopsis glomerata* | Cactaceae | Stem succulent | 0 |
| *Maihueniopsis glomerata* | Cactaceae | Stem succulent | 1 |
| *Malvastrum coromandelianum* | Malvaceae | Herbaceous non-graminoid | 1 |
| *Mandevilla laxa* | Apocynaceae | Climber | 1 |
| *Mandevilla pentlandiana* | Apocynaceae | Climber | 0 |
| *Manihot grahamii* | Euphorbiaceae | Shrub | 0 |
| *Margyricarpus pinnatus* | Rosaceae | Herbaceous non-graminoid | 1 |
| *Maytenus boaria* | Celastraceae | Tree | 0 |
| *Maytenus viscifolia* | Celastraceae | Tree | 0 |
| *Maytenus vitis-idaea* | Celastraceae | Shrub | 1 |
| *Melica macra* | Poaceae | Herbaceous graminoid | 1 |
| *Menodora integrifolia* | Oleaceae | Herbaceous non-graminoid | 0 |
| *Microchloa indica* | Poaceae | Herbaceous graminoid | 0 |
| *Mimozyganthus carinatus* | Fabaceae | Shrub | 1 |
| *Mitracarpus cuspidatus* | Rubiaceae | Herbaceous non-graminoid | 0 |
| *Monanthochloe littoralis* | Poaceae | Herbaceous graminoid | 1 |
| *Monteverdia spinosa* | Celastraceae | Shrub | 1 |
| *Monttea aphylla* | Plantaginaceae | Shrub | 0 |
| *Muhlenbergia peruviana* | Poaceae | Herbaceous graminoid | 1 |
| *Myrcianthes cisplatensis* | Myrtaceae | Tree | 1 |
| *Myriopteris myriophylla* | Pteridaceae | Fern | 0 |
| *Nassella filiculmis* | Poaceae | Herbaceous graminoid | 1 |
| *Nassella neesiana* | Poaceae | Herbaceous graminoid | 1 |
| *Nassella nidulans* | Poaceae | Herbaceous graminoid | 0 |
| *Nassella tenuissima* | Poaceae | Herbaceous graminoid | 1 |
| *Nassella trichotoma* | Poaceae | Herbaceous graminoid | 1 |
| *Neobouteloua lophostachya* | Poaceae | Herbaceous graminoid | 1 |
| *Nicotiana glauca* | Solanaceae | Shrub | 1 |

| | | | |
|---|---|---|---|
| *Nicotiana noctiflora* | Solanaceae | Herbaceous non-graminoid | 0 |
| *Nierembergia linariaefolia* | Solanaceae | Herbaceous non-graminoid | 1 |
| *Nothoscordum gracile* | Amaryllidaceae | Herbaceous graminoid | 1 |
| *Noticastrum argenteum* | Asteraceae | Herbaceous non-graminoid | 0 |
| *Noticastrum marginatum* | Asteraceae | Herbaceous non-graminoid | 1 |
| *Oenothera indecora* | Onagraceae | Herbaceous non-graminoid | 1 |
| *Oplismenus hirtellus* | Poaceae | Herbaceous graminoid | 1 |
| *Opuntia sulphurea* | Cactaceae | Stem succulent | 1 |
| *Oxalis conorrhiza* | Oxalidaceae | Herbaceous non-graminoid | 1 |
| *Pappophorum caespitosum* | Poaceae | Herbaceous graminoid | 1 |
| *Pappophorum malacophylum* | Poaceae | Herbaceous graminoid | 0 |
| *Pappophorum philippianum* | Poaceae | Herbaceous graminoid | 1 |
| *Parkinsonia praecox* | Caesalpiniaceae | Tree | 1 |
| *Parthenium hysterophorus* | Asteraceae | Herbaceous non-graminoid | 1 |
| *Pascalia glauca* | Asteraceae | Herbaceous non-graminoid | 0 |
| *Paspalum dilatatum* | Poaceae | Herbaceous graminoid | 0 |
| *Paspalum malacophyllum* | Poaceae | Herbaceous graminoid | 0 |
| *Paspalum notatum* | Poaceae | Herbaceous graminoid | 1 |
| *Paspalum quadrifarium* | Poaceae | Herbaceous graminoid | 1 |
| *Passiflora caerulea* | Passifloraceae | Climber | 1 |
| *Pavonia aurigloba* | Malvaceae | Herbaceous non-graminoid | 1 |
| *Pellaea ternifolia* | Pteridaceae | Fern | 0 |
| *Petiveria alliacea* | Petiveriaceae | Herbaceous non-graminoid | 0 |
| *Petunia axillaris* | Solanaceae | Herbaceous non-graminoid | 1 |
| *Pfaffia gnaphalioides* | Amaranthaceae | Herbaceous non-graminoid | 0 |
| *Phlegmariurus saururus* | Lycopodiaceae | Fern | 0 |
| *Piptochaetium montevidense* | Poaceae | Herbaceous graminoid | 1 |
| *Pitraea cuneato-ovata* | Verbenaceae | Herbaceous non-graminoid | 0 |
| *Plantago argentina* | Plantaginaceae | Herbaceous non-graminoid | 0 |
| *Plantago australis* | Plantaginaceae | Herbaceous non-graminoid | 0 |
| *Plantago brasiliensis* | Plantaginaceae | Herbaceous non-graminoid | 1 |
| *Plectrocarpa tetracantha* | Zygophyllaceae | Shrub | 1 |
| *Pleopeltis pinnatifida* | Polypodiaceae | Fern | 1 |
| *Poa ligularis* | Poaceae | Herbaceous graminoid | 0 |
| *Poa stuckertii* | Poaceae | Herbaceous graminoid | 1 |
| *Polylepis australis* | Rosaceae | Tree | 1 |
| *Polystichum montevidense* | Dryopteridaceae | Fern | 0 |
| *Polystichum pycnolepis* | Dryopteridaceae | Fern | 0 |
| *Porlieria microphylla* | Zygophyllaceae | Shrub | 1 |
| *Portulaca cryptopetala* | Portulacaceae | Herbaceous non-graminoid | 1 |
| *Portulaca grandiflora* | Portulacaceae | Herbaceous non-graminoid | 1 |
| *Portulaca oleracea* | Portulacaceae | Herbaceous non-graminoid | 1 |
| *Neltuma alba (Prosopis alba)* | Fabaceae | Tree | 1 |
| *Neltuma caldenia (Prosopis caldenia)* | Fabaceae | Tree | 1 |

| | | | |
|---|---|---|---|
| *Neltuma chilensis (Prosopis chilensis)* | Fabaceae | Tree | 1 |
| *Neltuma flexuosa (Prosopis flexuosa)* | Fabaceae | Tree | 1 |
| *Neltuma kuntzei (Prosopis kuntzei)* | Fabaceae | Tree | 1 |
| *Neltuma nigra (Prosopis nigra)* | Fabaceae | Tree | 1 |
| *Neltuma pugionata (Prosopis pugionata)* | Fabaceae | Tree | 0 |
| *Neltuma ruscifolia (Prosopis ruscifolia)* | Fabaceae | Tree | 0 |
| *Neltuma sericantha (Prosopis sericantha)* | Fabaceae | Shrub | 1 |
| *Strombocarpa strombulifera (Prosopis strombulifera)* | Fabaceae | Shrub | 1 |
| *Strombocarpa torquata (Prosopis torquata)* | Fabaceae | Shrub | 1 |
| *Pseudabutilon callimorphum* | Malvaceae | Shrub | 0 |
| *Pseudabutilon pedunculatum* | Malvaceae | Herbaceous non-graminoid | 1 |
| *Pseudognaphalium gaudichaudianum* | Asteraceae | Herbaceous non-graminoid | 1 |
| *Pteridium esculentum* | Dennstaedtiaceae | Fern | 0 |
| *Ranunculus praemorsus* | Ranunculaceae | Herbaceous non-graminoid | 0 |
| *Rhynchosia edulis* | Fabaceae | Herbaceous non-graminoid | 0 |
| *Rhynchosia senna* | Fabaceae | Herbaceous non-graminoid | 1 |
| *Ruprechtia apetala* | Polygonaceae | Tree | 1 |
| *Salicornia ambigua* | Amaranthaceae | Leaf succulent | 0 |
| *Salpichroa origanifolia* | Solanaceae | Herbaceous non-graminoid | 1 |
| *Salvinia nutans* | Salviniaceae | Herbaceous non-graminoid | 0 |
| *Sarcomphalus mistol* | Rhamnaceae | Tree | 1 |
| *Schinopsis haenkeana* | Anacardiaceae | Tree | 1 |
| *Schinopsis lorentzii* | Anacardiaceae | Tree | 1 |
| *Schinopsis quebracho-colorado* | Apocynaceae | Tree | 1 |
| *Schinus fasciculata* | Anacardiaceae | Shrub | 1 |
| *Schizachyrium condensatum* | Poaceae | Herbaceous graminoid | 1 |
| *Schizachyrium microstachyum* | Poaceae | Herbaceous graminoid | 1 |
| *Schizachyrium spicatum* | Poaceae | Herbaceous graminoid | 1 |
| *Schizachyrium tenerum* | Poaceae | Herbaceous graminoid | 0 |
| *Schkuhria pinnata* | Asteraceae | Herbaceous non-graminoid | 1 |
| *Selaginella sellowii* | Selaginellaceae | Herbaceous non-graminoid | 0 |
| *Senecio subulatus* | Asteraceae | Shrub | 0 |
| *Senegalia gilliesii* | Fabaceae | Shrub | 1 |
| *Senegalia praecox* | Fabaceae | Shrub | 1 |
| *Senna aphylla* | Fabaceae | Shrub | 1 |
| *Serpocaulon lasiopus* | Polypodiaceae | Fern | 0 |
| *Setaria hunzikeri* | Poaceae | Herbaceous graminoid | 1 |
| *Setaria lachnea* | Poaceae | Herbaceous graminoid | 0 |
| *Setaria oblongata* | Poaceae | Herbaceous graminoid | 0 |
| *Setaria pampeana* | Poaceae | Herbaceous graminoid | 1 |

| | | | |
|---|---|---|---|
| *Setaria parviflora* | Poaceae | Herbaceous graminoid | 1 |
| *Sida argentina* | Malvaceae | Herbaceous non-graminoid | 1 |
| *Sida dictyocarpa* | Malvaceae | Herbaceous non-graminoid | 1 |
| *Sida rhombifolia* | Malvaceae | Herbaceous non-graminoid | 1 |
| *Sida spinosa* | Malvaceae | Herbaceous non-graminoid | 0 |
| *Sida variegata* | Malvaceae | Herbaceous non-graminoid | 0 |
| *Sisyrinchium chilense* | Iridaceae | Herbaceous non-graminoid | 0 |
| *Sisyrinchium unguiculatum* | Iridaceae | Herbaceous graminoid | 1 |
| *Solanum amygdalifolium* | Solanaceae | Herbaceous non-graminoid | 0 |
| *Solanum angustifidum* | Solanaceae | Herbaceous non-graminoid | 0 |
| *Solanum argentinum* | Solanaceae | Shrub | 1 |
| *Solanum chacoense* | Solanaceae | Herbaceous non-graminoid | 0 |
| *Solanum elaeagnifolium* | Solanaceae | Herbaceous non-graminoid | 1 |
| *Solanum palinacanthum* | Solanaceae | Herbaceous non-graminoid | 0 |
| *Solanum pseudocapsicum* | Solanaceae | Herbaceous non-graminoid | 0 |
| *Solanum sisymbriifolium* | Solanaceae | Herbaceous non-graminoid | 0 |
| *Solanum tuberosum* | Solanaceae | Herbaceous non-graminoid | 0 |
| *Solidago chilensis* | Asteraceae | Herbaceous non-graminoid | 1 |
| *Sorghastrum pellitum* | Poaceae | Herbaceous graminoid | 1 |
| *Spergula ramosa* | Caryophyllaceae | Herbaceous non-graminoid | 0 |
| *Spermacoce eryngioides* | Rubiaceae | Herbaceous non-graminoid | 0 |
| *Sphaeralcea cordobensis* | Malvaceae | Herbaceous non-graminoid | 1 |
| *Sporobolus indicus* | Poaceae | Herbaceous graminoid | 1 |
| *Sporobolus pyramidatus* | Poaceae | Herbaceous graminoid | 1 |
| *Stenandrium dulce* | Acanthaceae | Herbaceous non-graminoid | 1 |
| *Stetsonia coryne* | Cactaceae | Stem succulent | 1 |
| *Stevia achalensis* | Asteraceae | Shrub | 0 |
| *Stevia satureifolia* | Asteraceae | Shrub | 0 |
| *Struthanthus uraguensis* | Loranthaceae | Herbaceous non-graminoid | 1 |
| *Stylosanthes gracilis* | Fabaceae | Herbaceous non-graminoid | 0 |
| *Stylosanthes guianensis* | Fabaceae | Herbaceous non-graminoid | 0 |
| *Suaeda divaricata* | Chenopodiaceae | Shrub | 1 |
| *Tagetes argentina* | Asteraceae | Herbaceous non-graminoid | 1 |
| *Tagetes filifolia* | Asteraceae | Herbaceous non-graminoid | 0 |
| *Tagetes minuta* | Asteraceae | Herbaceous non-graminoid | 1 |
| *Talinum fruticosum* | Talinaceae | Herbaceous non-graminoid | 0 |
| *Talinum paniculatum* | Talinaceae | Herbaceous non-graminoid | 1 |
| *Talinum polygaloides* | Talinaceae | Herbaceous non-graminoid | 1 |
| *Taraxacum officinale* | Asteraceae | Herbaceous non-graminoid | 1 |
| *Tephrocactus alexanderi* | Cactaceae | Stem succulent | 0 |
| *Tephrocactus articulatus* | Cactaceae | Stem succulent | 1 |
| *Thelypteris argentina* | Thelypteridaceae | Fern | 0 |
| *Tillandsia aizoides* | Bromeliaceae | Bromeliad | 1 |
| *Tillandsia bryoides* | Bromeliaceae | Bromeliad | 1 |

| | | | |
|---|---|---|---|
| *Tillandsia capillaris* | Bromeliaceae | Bromeliad | 1 |
| *Tillandsia duratii* | Bromeliaceae | Bromeliad | 1 |
| *Tillandsia myosura* | Bromeliaceae | Bromeliad | 1 |
| *Tillandsia rectangula* | Bromeliaceae | Bromeliad | 1 |
| *Tillandsia usneoides* | Bromeliaceae | Bromeliad | 1 |
| *Tillandsia xiphioides* | Bromeliaceae | Bromeliad | 1 |
| *Tragia hieronymi* | Euphorbiaceae | Herbaceous non-graminoid | 1 |
| *Tribulus terrestris* | Zygophyllaceae | Herbaceous non-graminoid | 1 |
| *Trichocereus candicans* | Cactaceae | Stem succulent | 0 |
| *Tricomaria usillo* | Malpighiaceae | Shrub | 1 |
| *Trifolium amabile* | Fabaceae | Herbaceous non-graminoid | 0 |
| *Trifolium repens* | Fabaceae | Herbaceous non-graminoid | 1 |
| *Triglochin scilloides* | Juncaginaceae | Herbaceous non-graminoid | 0 |
| *Tripogonella spicata* | Poaceae | Herbaceous graminoid | 0 |
| *Trithrinax campestris* | Arecaceae | Tree | 1 |
| *Trixis divaricata* | Asteraceae | Herbaceous non-graminoid | 0 |
| *Troncosoa seriphioides* | Verbenaceae | Shrub | 0 |
| *Turnera sidoides* | Turneraceae | Herbaceous non-graminoid | 1 |
| *Vachellia aroma* | Fabaceae | Shrub | 1 |
| *Vachellia caven* | Fabaceae | Shrub | 1 |
| *Valeriana ferax* | Valerianaceae | Herbaceous non-graminoid | 0 |
| *Verbena litoralis* | Verbenaceae | Herbaceous non-graminoid | 0 |
| *Verbesina encelioides* | Asteraceae | Herbaceous non-graminoid | 1 |
| *Vernonanthura nudiflora* | Asteraceae | Shrub | 1 |
| *Vicia graminea* | Fabaceae | Herbaceous non-graminoid | 1 |
| *Woodsia montevidensis* | Woodsiaceae | Fern | 0 |
| *Xanthium spinosum* | Asteraceae | Herbaceous non-graminoid | 0 |
| *Xanthium strumarium* | Asteraceae | Herbaceous non-graminoid | 0 |
| *Ximenia americana* | Olacaceae | Tree | 1 |
| *Zanthoxylum coco* | Rutaceae | Tree | 1 |
| *Zephyranthes longistyla* | Amaryllidaceae | Herbaceous non-graminoid | 0 |
| *Zinnia peruviana* | Asteraceae | Herbaceous non-graminoid | 1 |
| *Zuccagnia punctata* | Fabaceae | Shrub | 1 |

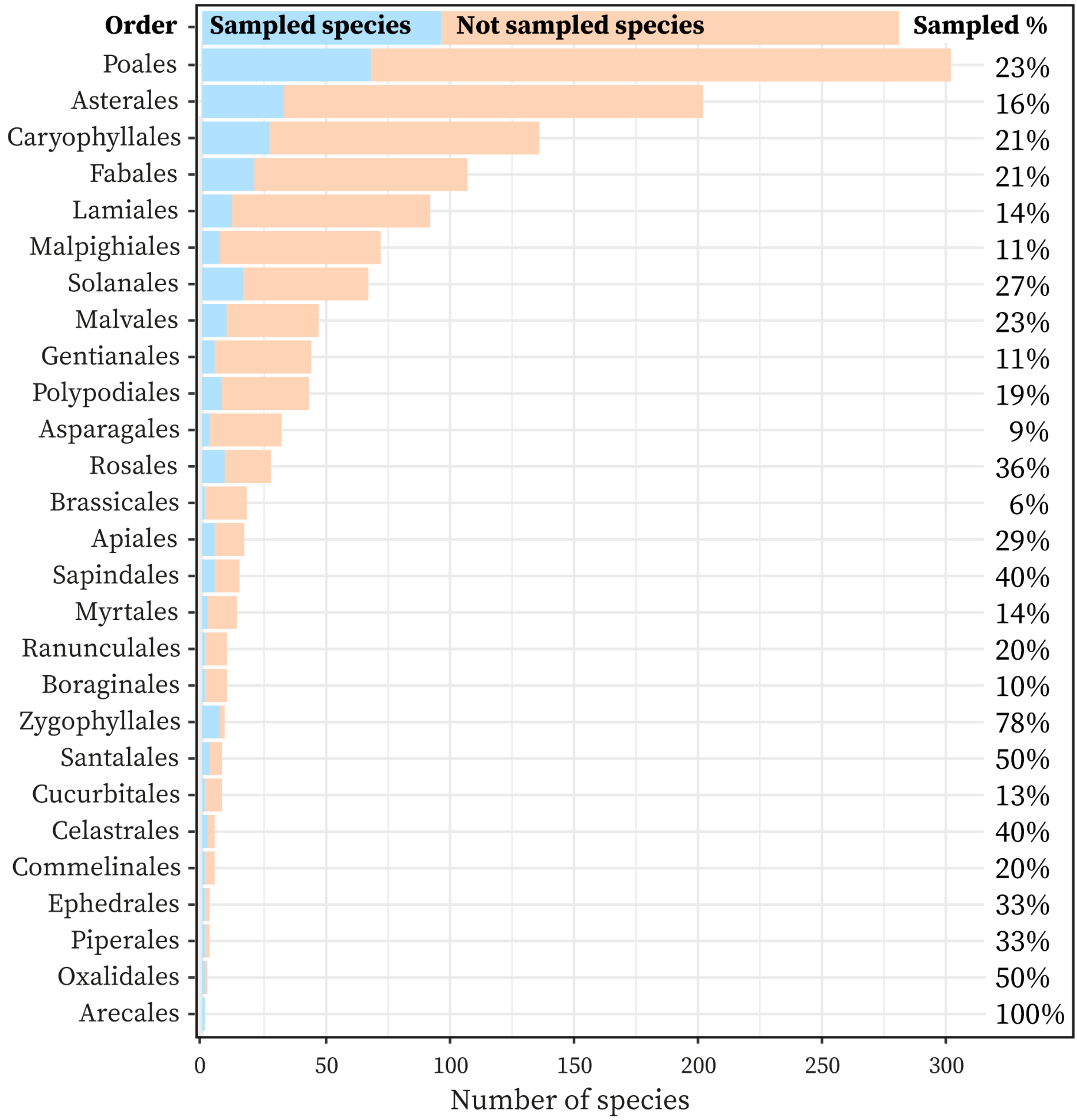


**FIGURE S1** Number of species of the flora of Central Argentina within different taxonomic orders considered in this study. The blue bars indicate the number of species on which all six traits of the GSPFF were measured, and the pink ones, the number of species in each order previously recorded in comprehensive surveys of the flora of Central Argentina (Cabido et al., 1994; Cantero et al., 2001; Giorgis et al., 2017) that were not sampled in the present study. The sum of both bars indicates the total number of species documented in the regional flora for each taxonomic group (the 100% out of which the percentage sampled was calculated, indicated to the right of each bar). There are another 13 plant orders present in the Central Argentina flora, containing 7 or fewer species each, that were not sampled by us and are not represented in the figure (Geraniales with 7 species; Ericales and Cornales with 6; Dipsacales, Ophioglossales and Saxifragales with 4; Selaginellales with 3; Alismatales and Lycopodiales with 2; and Equisetales, Escalloniales, Hymenophyllales and Vitales with 1).

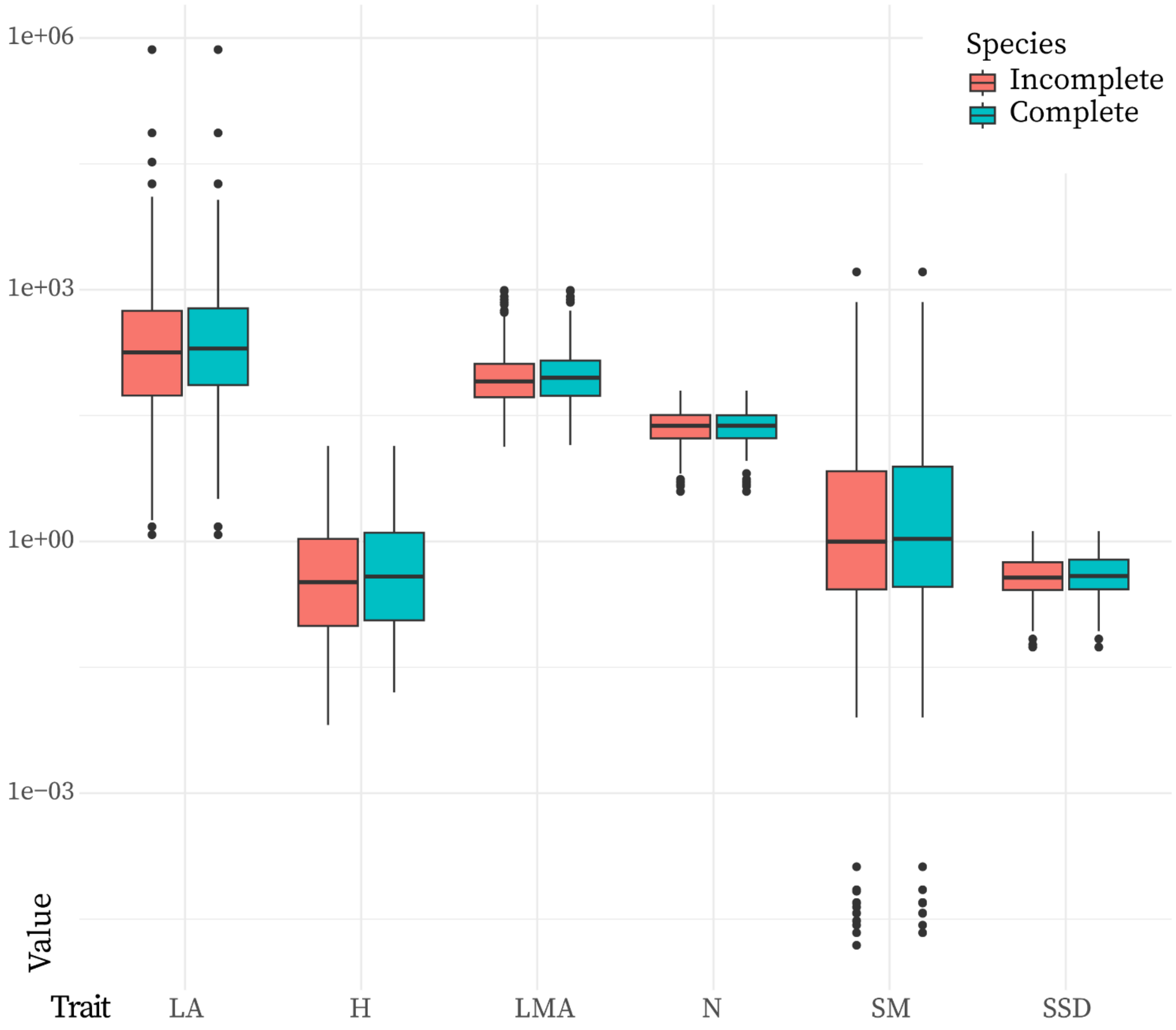


**FIGURE S2** Comparison between species with incomplete and complete functional trait measurements. The boxplots represent the distributions of the six functional traits for the species with complete (268) and incomplete (410) empirical measurements of the six traits.

**TABLE S3** Bivariate relationships in the regional and global flora between the six functional traits analyzed. The table shows the standardized major axis statistics, slope with its statistical significance and $R^2$ (Warton et al., 2006). The significance levels are the following: (*) 0.05 > p > 0.01; (**) 0.01 > p > 0.001; (***) p < 0.001.

| | Slope | | $R^2$ | |
|---|---|---|---|---|
| Trait combination | Global | Regional | Global | Regional |
| H-SM | 1.92*** | 1.87*** | 0.31 | 0.16 |
| H-SSD | 0.28*** | 0.37*** | 0.59 | 0.15 |
| H-LA | 1.06*** | 1.20*** | 0.34 | 0.08 |
| H-LMA | 0.32*** | 0.47*** | 0.08 | 0.07 |
| H-N | -0.21*** | 0.30* | 0.01 | 0.02 |
| SM-SSD | 0.18*** | 0.20*** | 0.32 | 0.09 |
| SM-LA | 0.58*** | 0.66 | 0.15 | 0.01 |
| SM-LMA | 0.19*** | 0.25*** | 0.06 | 0.08 |
| SM-N | -0.13 | 0.16*** | 0.00 | 0.04 |
| SSD-LA | 3.37*** | -3.13*** | 0.10 | 0.11 |
| SSD-LMA | 0.99*** | 1.25* | 0.36 | 0.02 |
| SSD-N | -0.67*** | 0.78 | 0.08 | 0.01 |
| LA-LMA | -0.29*** | -0.39 | 0.01 | 0.00 |
| LA-N | 0.21*** | 0.25* | 0.02 | 0.01 |
| LMA-N | -0.70*** | -0.61*** | 0.34 | 0.24 |

**TABLE S4** Principal component analyses (PCA) of the regional flora of Central Argentina and the global flora as represented by the GSPFF (Díaz et al., 2016) based on six functional traits. The table shows the percentage of variation explained and eigenvalues for the first two principal components (PC1 and PC2) for the regional PCA and the GSPFF (2016). Below are the loading and rank of each trait for PC1 and PC2, as well as the ranking of loadings.

| | **Regional PC1** | | **Global PC1** | | **Regional PC2** | | **Global PC2** | |
|---|---|---|---|---|---|---|---|---|
| Variation explained (%) | 32.82 | | 48.88 | | 25.30 | | 24.96 | |
| Eigenvalue | 1.97 | | 2.93 | | 1.52 | | 1.50 | |
| **Trait** | **Rank** | **Load** | **Rank** | **Load** | **Rank** | **Load** | **Rank** | **Load** |
| Adult plant height (H) | 1 | -0.59 | 1 | 0.52 | 4 | -0.14 | 5 | 0.20 |
| Diaspore mass (SM) | 2 | -0.55 | 3 | 0.45 | 5 | -0.10 | 4 | 0.30 |
| Stem specific density (SSD) | 3 | -0.49 | 2 | 0.51 | 6 | 0.05 | 6 | -0.09 |
| Leaf area (LA) | 6 | -0.01 | 6 | 0.23 | 3 | -0.32 | 1 | 0.58 |
| Leaf mass per area (LMA) | 4 | -0.32 | 4 | 0.40 | 2 | 0.60 | 3 | -0.46 |
| N content per unit leaf mass (Nmass) | 5 | -0.10 | 5 | -0.25 | 1 | -0.71 | 2 | 0.57 |
